# Evolution of Crassulacean Acid Metabolism (CAM) as an escape from ecological niche conservatism in Malagasy *Bulbophyllum* (Orchidaceae)

**DOI:** 10.1101/2021.01.09.426039

**Authors:** Alexander Gamisch, Winter Klaus, Gunter A. Fischer, Hans Peter Comes

## Abstract

Despite growing evidence that niche shifts are more common in flowering plants than previously thought, still little is known about the key physiological (e.g. photosynthesis) traits underlying such niche shifts. To address this question, we here combine a comprehensively sampled phylogeny for mostly epiphytic Malagasy *Bulbophyllum* orchids (*c*. 210 spp.) with climatic niche and carbon isotope-derived photosynthesis data to infer the groups’ spatial-temporal history and to test the role of Crassulacean Acid Metabolism (CAM), a highly water-use efficient type of photosynthesis, in facilitating niche shifts and diversification. We find that most extant species still retain niche (Central Highland) and photosynthesis (C_3_) states as likely present in the single mid-Miocene (*c.* 12.70 Ma) ancestor colonizing Madagascar. However, we also infer a major transition to CAM, linked to a late Miocene (*c*. 7.36 Ma) invasion of species from the sub-humid highland niche first into the island’s humid eastern coastal, and then into the seasonally dry ‘Northwest Sambirano’ rainforests, yet without significant effect on diversification rates. These findings support the rarely recognized hypothesis that CAM in tropical epiphytes may be selectively advantageous even in high rainfall habitats, rather than presenting a mere adaptation to dry environments or epiphytism *per se*. Overall, our study qualifies CAM as an evolutionary ‘gateway’ trait that considerably widened the spatial-ecological amplitude of Madagascar’s most species-rich orchid genus.

## Introduction

Analysing the extent to which species or clades are able to retain or shift their environmental niche space over time is key to understanding large-scale patterns of lineage diversification and adaptations to environmental change. The retention of niche-related ancestral ecological traits over time has been termed niche conservatism (Wiens *et al.* 2010), while evolutionary change that allows lineages to transverse niche boundaries is known as niche shift (Donoghue & Edwards 2014). Niche conservatism has long been viewed as the dominant process in ecology across many flowering plant (angiosperm) lineages (Crisp *et al*. 2009; Donoghue & Edwards 2014), but there is growing evidence that niche shifts are more common than previously thought (Holstein & Renner 2011; Evans *et al*. 2014; Gamisch *et al.* 2016; Cardillo *et al.* 2017; Herrera 2020; Dale *et al.* 2020). However, still little is known about the key structural and physiological traits underlying such niche shifts, and their consequences for species diversification at broad phylogenetic and spatial scales (reviewed in Soltis *et al*. 2019).

In many groups of angiosperms, niche shifts tend to occur from moister to drier habitats and often involve adaptations to higher light exposure and/or lower water supply, such as a succulent morphology and water-saving forms of photosynthesis such as C_4_ and especially CAM photosynthesis (Evans *et al.* 2014; Donoghue & Edwards 2014; Bone *et al.* 2015; Males 2017). The water-conserving CAM pathway of photosynthesis (crassulacean acid metabolism) has been documented in 37 mostly angiosperm families thus far (Winter *et al*. 2021), and has evolved independently from C_3_ ancestors many times, even within genera (Winter *et al.* 1983; Winter & Smith 1996; Crayn *et al.* 2004; Silvera *et al.* 2010, 2014; Bone *et al.* 2015; Heyduk *et al.* 2019; Li *et al.* 2019). In CAM plants, overall transpirational water loss is reduced owing to stomatal closure for most of the day when the driving forces for water loss are high. Stomata open predominantly at night concomitant with nocturnal uptake of CO_2_ when the driving forces for water loss are low (Winter & Smith 1996; Lüttge 2004; Heyduk *et al.* 2019). CAM occurs in many succulent species of warm, semi-arid regions (e.g. Cactaceae), but also in many tropical vascular epiphytes that are subject to enormous short-term and seasonal variations in the availability of water (Keddy 2007). CAM is well-known in tropical epiphytic species of Bromeliaceae and Orchidaceae, where CAM is generally considered a ‘key innovation’, promoting the invasion of new habitats and accelerating species diversification (Gravendeel *et al*. 2004; Motomura *et al.* 2008; Silvestro *et al*. 2013; Bouchenak-Khelladi *et al*. 2015; Givnish *et al.* 2014, 2015). However, strong evidence in support of this notion is hard to come by as ‘key innovations’ are difficult to test. For example, diversification rates were not significantly different between CAM and C_3_ species of Afro-Malagasy Eulophiinae orchids (Bone *et al.* 2015). Moreover, in an Asian clade of the pantropical orchid genus *Bulbophyllum* Thouars, CAM species actually suffered from a tenfold higher extinction rate compared to their C_3_ con-generics (Hu 2018). Thus, there is still a big gap in our knowledge regarding the extent to which CAM is linked to diversification and niche evolution in Orchidaceae, where less than 10% of all species (*c*. 26,000 in total; WCSP 2020) have so far been surveyed for mode of photosynthesis (Silvera *et al.* 2009, 2010a,b; Winter 2019; Torres-Morales *et al.,* 2020).

Here, we study a monophyletic and mostly epiphytic (or rarely litho-/rheophytic) lineage of *Bulbophyllum* from Madagascar (*c*. 210 spp.; Sieder *et al.* 2009; Madagascar Catalogue 2019) to further clarify the extent to which CAM is linked to niche evolution and species diversification in tropical orchids. Based on a genus-wide phylogeny of nuclear ribosomal (nr) DNA (Gamisch & Comes 2019), this hyper-diverse island radiation of *c*. 210 species likely dates back to the mid-Miocene (*c.* 12.36 million years ago, Ma). Species richness of this lineage is highest in mid-elevation sub-humid forest (*c*. 800–1,300 m) along the eastern slopes of the ‘Central Highlands’ (Cornet 1974; Yoder & Nowak 2006; Fischer *et al.* 2007; Gamisch *et al.* 2016). However, members of a small subclade (termed ‘C’, *c*. 30 spp.; Gamisch *et al.* 2015, 2016) also occur in eastern-coastal, warm-humid evergreen rainforest and/or seasonally dry ‘Northwest Sambirano’ rainforest (both *c*. 0–800 m; Gamisch *et al*. 2016), possibly reflecting major niche and/or physiological transitions during the history of Malagasy *Bulbophyllum*. However, in *Bulbophyllum*, a limited survey of carbon isotope ratios (^13^C/^12^C) has so far revealed only C_3_ photosynthesis in Madagascar (Kluge *et al*. 1995), while Austral-Asian species are known to exhibit both C_3_-and CAM-type isotopic signatures (Winter *et al.* 1983; Earnshaw *et al.* 1987; Hu 2018; see above).

Here, we report the most comprehensive (nuclear/plastid) phylogenetic and carbon isotope datasets for Malagasy *Bulbophyllum* to date, covering *c*. 85.2% and 90.5% of the total species diversity, respectively. Based on this, we then combine diversification rate analyses with ancestral climatic niche and photosynthetic pathway reconstructions to (1) infer the group’s spatial-temporal history; and (2) to test the role of CAM in facilitating niche shifts and diversification in Madagascar’s most species-rich orchid lineage. Moreover, we test hypotheses of broader biogeographical and evolutionary-physiological significance, namely that (3) insular radiations typically begin in marginal (coastal) rather than interior habitats (Liebherr & Hajek 1990; Schluter 2000); and that (4) CAM in tropical epiphytes may be selectively advantageous even in high rainfall forests (Pierce *et al.* 2002; Lüttge 2004), rather than presenting a mere adaptation to dry environments or epiphytism *per se* (e.g. Gravendeel *et al.* 2004; Silvera *et al.* 2009, 2010; Bone *et al.* 2015).

## Materials and Methods

### Phylogenetic taxon sampling and molecular dating

We inferred phylogenetic relationships among 179 species of Malagasy *Bulbophyllum* [*c*. 85.2% of the group’s total (210 spp.) diversity; Sieder *et al.* 2009], using DNA sequence information from three nuclear (nrITS, *PI, Xdh*) and five plastid gene regions (*atp*I–*atp*H, *ycf1*, *mat*K, *trn*D*–trn*E, *psb*A*–trn*H; see Supplemental **Table S1**). This taxon sampling includes 22 undetermined and five newly described species (Fischer *et al.* 2009; Hermans *et al*., in press), plus nine species from Asia (four spp.), Africa (three) and the Neotropics (two) (Supplemental **Table S1**). Sequences were obtained from GenBank (Fischer *et al.* 2007; Gamisch *et al.* 2015; Gamisch & Comes 2019) or newly generated (*Xdh, mat*K) following Gorniak *et al.* (2010) and Ford *et al.* (2009), aligned by ClustalW in GENEIOUS v.10.2.3 (Kearse *et al.* 2012) and manually edited. The total alignment of the 188 species (7712 bp) included 1504 sequences, *c*. 87% (1306) of which were complete (Supplemental **Table S1**). Of the 7712 nucleotide sites, *c.* 15.4% (1191) were parsimony informative (**Table S2**).

All plastid regions were combined and treated as a single partition, while the three nuclear regions were analysed separately. For each partition, the best fitting model of nucleotide substitution was identified (**Table S2**) using the size-corrected Akaike Information Criterion (AICc) in IQ-TREE v.1.6.2 (http://iqtree.cibiv.univie.ac.at; Trifinopoulos *et al.* 2016). Phylogenetic species relationships and divergence times were estimated in BEAST v.1.8.4 (Drummond *et al.* 2012) using age estimates of a previous genus-wide (nrITS) phylogeny (Gamisch & Comes 2019) as secondary calibration points: (1) a uniform prior distribution ranging from 16.2 to 25.63 Ma at the *Bulbophyllum* root; (2) a normal prior of 12.26 ± 2.28 Ma (mean ± standard deviation, SD) for the crown of Malagasy *Bulbophyllum*; and (3) a normal prior of 5.67 ± 1.31 Ma for the crown of subclade C. A relaxed molecular clock with uncorrelated log-normal model was used (Drummond *et al.* 2006). The tree speciation prior followed a constant-rates birth-death (CR-BD) process, and one Markov chain Monte Carlo (MCMC) run was performed on the CIPRES Science Gateway portal (Miller *et al.* 2010) for 2 × 10^8^ generations, sampling every 20,000^th^ step. TRACER v.1.7.1 (Rambaut *et al.* 2018) was used to confirm that all parameters had large enough effective sample sizes (ESS > 233) after discarding the first 10% of sampled tress as burn-in.

### Assembly of georeferenced presence records and climatic niche datasets

In a first step, we assembled 5116 georeferenced presence records of Malagasy *Bulbophyllum* (coordinates available upon request), representing all named species plus 895 records of unknown identity. Records were sourced from own expeditions (2000–2018) in Madagascar (G.A.F., A.G. and Anton Sieder, Vienna) and La Réunion (A.G. and Ursula Jaros), the Global Biodiversity Information Facility (GBIF Occurrence Download http://doi.org/10.15468/dl.ehvjtf; 7th July 2017), and voucher specimens obtained from herbaria (G, K, MO, NEU, P, SZU, TAN, W, WU). Duplicate presence records per species were discarded. In a second step, we used SPTHIN v.0.1.0 (Aiello-Lammens *et al.* 2015) to generate two spatially filtered datasets (Merow *et al.* 2013): in *dataset 1*, all records within a 1-km radius were pruned to one random accession, regardless of species identity (*n* = 1206); in *dataset 2*, all records per species within such a distance were pruned to one random accession (*n* = 3036). Hence, *dataset 1* adequately represents the climatic niche space of Malagasy *Bulbophyllum* at genus- vs. species-level, as best suited for traditional (qualitative) ancestral state reconstructions, while *dataset 2* provides continuous, interspecific trait data as required for comparative phylogenetic methods (PCMs; see below). For each locality and dataset, we extracted 19 bioclimatic variables (bio1–19) from WORLDCLIM v.1.4 (Hijmans *et al.* 2005) at 30 arc-seconds resolution for current conditions (*c.* 1950–2000). Note, four species accessions for which no precise locality data were available were not included in *dataset 2* and thus pruned from subsequent analyses.

### Identification of macroecological niche states

For identifying macroecological niche states, we first subjected *dataset 1* (1206 locality-climate points) to principal components analysis (PCA) in R v.3.6.1. (R Core Team 2019) To complement traditional PCA loadings, we also calculated *R*^2^ values for each climate variable and ordinated axis (PC1, PC2) through regression of mean trait values per species on their respective mean PC scores to estimate which variables were most important (Folk *et al.* 2019). To investigate how these data points group in multi-dimensional climatic space, we used *k*-means clustering, as selected by CLVALID v.0.6-6 (Brock *et al.* 2011) when tested against alternative algorithms (e.g. hierarchical clustering, partitioning around medoids; results not shown). Accordingly, we used the *kmeans* function in NBCLUST v.3.0 (Charrad *et al.* 2014) to determine the optimal number of clusters (*K*) in this dataset based on 15 validity indices (*kl, ch, hartigan, cindex, db, silhouette, ratkowsky, ball, ptbiserial, gap, frey, mcclain, dunn, sdindex, sdbw*). Clustering and cluster visualization were carried out in CLUSTER v.2.0.6 (Maechler 2017) and FACTOEXTRA v.1.0.5 (Kassambara & Mund 2017). This analysis identified three clusters (viz. niches), hereafter referred to as *A*, *B*, and *C* (see Results).

For each niche, we generated ecological niche models (ENMs) in MAXENT v.3.4.1 (Phillips *et al.* 2017) using ten bioclimatic layers (bio1, 3, 4, 6, 7, 12, 16–19), i.e. after removal of highly correlated variables (Pearson’s *r* ≥ |0.9|) in ENMTOOLS v.1.4.4 (Warren *et al.* 2010) to avoid potential overfitting. Fifty bootstrap replicates and 75% of the localities were used to train the models. We averaged the area under the Receiver Operating Characteristic (ROC) Curve (AUC) across replicates to determine the predictive power of each model (AUC > 0.7; Fawcett 2006). Suitable areas were defined as areas with higher occurrence (cloglog) probabilities than the ‘maximum training sensitivity plus specificity’ (MTSS) threshold (Liu *et al.* 2013). Niche range overlap was calculated with ENMTOOLS.

### Reconstruction of ancestral niche states and estimation of transition rates

Ancestral niche states were reconstructed on the BEAST-derived maximum clade credibility (MCC) chronogram of Malagasy *Bulbophyllum* (with outgroups pruned prior to analysis) using the submodule MULTISTATE of BAYESTRAITS v.3.0 (Pagel *et al.* 2004). For each species, niche state (*A*, *B* and/or *C*) was coded according to the mean ENM cluster suitability (above the MTSS) of its locality data (based on *dataset 2*). For instance, when the mean suitability of a given species was higher than the MTSS for *A* then this species was coded as *A*. If instead a species had similar mean suitability above the MTSS for two or more states (i.e. due to outliers and/or overlapping suitability in transitional zones between niches), a decision was made based on visual inspection of the species’ localities; otherwise, it was coded as ‘polymorphic’ (e.g. *BC* or *AB*). The MCMC analyses were run for 5 x 10^6^ generations, using a reversible-jump (rj) hyper-prior with an exponential prior (mean seeded from a uniform distribution on the interval 0 to100), and a burn-in of 5 x 10^4^ generations. The reconstructed niche states were plotted for nodes of interest, i.e. the crown node of Malagasy *Bulbophyllum*, the crown nodes of major clades, and all nodes of the ecologically diverse subclade C.

In addition, we used BAYESTRAITS to infer the direction of niche state transitions (*q*) by comparing a full (six-parameter) model with unconstrained parameters (i.e. *q*_AB_*, q*_AC_*, q*_BA_, *q*_BC_, *q*_CA_*, q*_CB_) to six models of zero uni-directional (i.e. asymmetrical) transitions for all pairwise combinations of states (i.e. *q*_AB_ = 0; *q*_AC_ = 0;…*q*_CB_ = 0) (see **Table S3**). Model selection followed 2 logarithmic Bayes Factors (BFs), with values > 6 (or 10) considered as strong (or very strong) evidence for one model over the other (Kass & Raftery 1995).

### Reconstruction of ancestral photosynthetic pathways (C3 vs. CAM)

Nocturnal fixation of atmospheric CO_2_ into malic acid is characteristic of CAM photosynthesis and results into a higher accumulation of ^13^C in plant carbon when compared to C_3_ photosynthesis (O’Leary 1988; Bone *et al.* 2015). Thus, carbon isotope ratios (^13^C/^12^C) can be used as proxy for photosynthetic pathway (Osmond *et al.* 1973). Accordingly, we performed leaf carbon isotope measurements in a total of 201 *Bulbophyllum* accessions, representing 192 different species (190 from Madagascar plus two outgroup species), 171 of which were included in the phylogeny of the present study (see also **Table S4**). Leaf samples were collected and silica-dried in the field (2000–2018) or obtained from herbarium specimens (WU, SZU, MO, P). About 2 mg per sample were subjected to carbon stable isotopic composition analyses (δ13C) using the Thermo Scientific Delta V Advantage isotope ratio mass spectrometer (IRMS) at the Smithsonian Tropical Research Institute (STRI), Balboa/Panama City, Panama. δ13C values were calculated relative to the Pee Dee Belemnite (PDB) standard (*Belemnitella americana*) using the formula: δ13C (‰) = [(^13^C/^12^C sample) / (^13^C/^12^C standard) − 1] × 1000 (Silvera *et al.* 2009). Following Crayn *et al.* (2004) and Winter *et al.* (2015), species with δ13C values less or more negative than −20‰ were binary coded as (strong) CAM or C_3_, respectively (cf. Bone *et al.* 2015; Li *et al.* 2019; Winter 2019). While values less negative than −20‰ indicate CO_2_ fixation predominantly via CAM, values more negative than −20‰ do not preclude the possibility of low-level CAM, but indicate that this did not make a major contribution to carbon gain. Ancestral character states for C_3_ vs. CAM were reconstructed in Bayestraits, using the same methods and settings as described above for ancestral niche states. Further, we performed separated runs in the Bayestraits submodule DISCRETE to test for independent vs. dependent (correlated) evolution of niche and photosynthetic states (Pagel & Meade 2006). As this test requires binary traits, niche *A* was subsumed together with *B*, resulting in a niche coding *C* vs. not *C* (or *C* vs. *AB*). Model selection followed BFs (see above).

### Estimating rates of niche evolution and lineage diversification

For quantifying rates of niche evolution, we first subjected *dataset 2* (3036 locality-climate points) to PCA (as described above) and calculated mean scores per species based on the first two components (PC1, PC2), respectively. We then used BAMM v.2.5.0 (Rabosky 2014) to infer rates of niche evolution along the MCC chronogram (with outgroups pruned) by performing a run with 10^6^ generations for each component (PC1, PC2) separately. Similarly, we used BAMM to estimate rates of net lineage diversification [*r* = speciation (λ) – extinction (*μ*)], taking incomplete taxon sampling into account (sampling frequency = 0.852). One rjMCMC run was performed with 10^7^ generations, sampling every 1000^th^ step under the default prior assumption of a single expected rate shift. Initial priors were obtained using the *setBAMMpriors* function in BAMMTOOLS v.2.1.6 (Rabosky *et al.* 2014). Post-analysis and visualization of rates through time were done in BAMMTOOLS after a 10% burn-in (see above).

In addition, we fitted ten birth-death (BD) models to the MCC chronogram with either time-dependent (six models), palaeo-temperature-dependent (two) or diversity-dependent (DD) diversification rates (two; see **Table S5**), using, respectively, the *fit_bd* and *fit_env* functions of RPANDA v.1.7 (Morlon *et al.* 2016) and the *dd_ML* function of DDD v.3.7 (Etienne *et al.* 2012), with each analysis accounting for unsampled taxa (see above). Exponential diversification dependencies were specified (in *fit_bd* and *fit_env*) since linear diversification dependencies can lead to unrealistic model fits (Gamisch 2020). The palaeo-temperature time curve employed (Hansen *et al.* 2013) was smoothed using the default ‘NULL’ option for the degree of freedom (d.f.), resulting in a d.f. of 100.06 palaeo-temperature. Models were considered superior if ΔAICc was > 2.0 (Burnham & Anderson 2002). Note, all the above analyses (Bamm; Rpanda/Ddd) were also run separately on subclade C and the pruned tree without this lineage.

Finally, we used the STRAPP (Structured Rate Permutations on Phylogenies) procedure in Bammtools (Rabosky & Huang 2016) to test for a significant relationship between mean *r* at the tips and the corresponding extant (1) quantitative niche states (PC1, PC2); or (2) the three macro-ecological niche states identified (*A*–*C*; see Results). Statistical significance was calculated using Pearson’s correlation coefficient and Kruskal-Wallis test, respectively, each based on 1000 permutations (two□tailed).

## Results

### Phylogeny reconstruction and divergence time estimates

Our BEAST-derived phylogeny based on three nuclear and five plastid gene regions (**Fig. S1**) supported the monophyly of Malagasy *Bulbophyllum* (posterior probability, PP = 1.0) and identified three main clades (termed A, B, and D), which essentially formed an unresolved trichotomy. Clades A (PP = 0.91) and B (PP = 1.0) harboured the great majority of species diversity (50 and 126 spp., respectively), while the three species of sect. *Inversiflora* (*sensu* Hermans *et al*., in press) formed clade D (PP = 1.0). Notably, subclade C (*sensu* Gamisch *et al.* 2015, 2016), comprised of sects. *Bifalcula* + *Calamaria* (30 spp.; PP = 1.0), occupied a nested and thus derived position within clade A, as likely sister of sect. *Kainochilus* (PP = 0.85; **Fig. S1**). Based on our molecular dating, we estimated the mean crown age of Malagasy *Bulbophyllum* to the mid-Miocene (*c*. 12.70 Ma; 95% highest posterior density, HPD: 10.39–15.27 Ma). Clades A and B started to diversify shortly thereafter in quick succession at *c*. 11.58 (9.16–13.87) Ma and 10.92 (8.68–13.14) Ma, respectively. In contrast, subclade C began to radiate in the late Miocene, *c*. 7.34 (5.83–8.89) Ma. The great majority of extant species (123/179, *c.* 71.1%) likely originated during the Quaternary (≤ 2.6 Ma; see **Fig. S1**).

### Identification of macroecological niche states

For the 1206 locality-climate data of Malagasy *Bulbophyllum* (*dataset 1*), five (out of 15) *k*-means validity indices each suggested clustering of the samples into two or three groups, respectively (data not shown). However, we favoured *K* = 3 as the optimal (i.e. more informative) solution as this partition recovered essentially the same clusters (*A*, *B*, *C*) as previously inferred for subclade C (Gamisch *et al.* 2016). In the corresponding PCA biplot (**Fig. S2**), samples of clusters *C* (*n* = 731) and *B* (*n* = 341) mainly differed along PC1 (accounting for 55.9% of the total variance), which was strongly (*R*^2^ > 0.7) and positively correlated with measures of temperature (bio1, 6, 8–11) and annual precipitation (bio12) (Supplemental **Table S6**). By contrast, cluster *A* (*n* = 134) differed along PC2 (22%), which had relatively strong (*R*^2^ > 0.4) and positive loadings of temperature seasonality (bio4) but negative ones of both isothermality (bio3) and precipitation seasonality (bio15) (**Table S6**). All pairwise comparisons between clusters revealed significant differences for each of the two axes tested (Mann-Whitney test, all *P* < 0.001).

The present-day ENMs generated for clusters *A*, *B*, and *C* (hereafter ‘niches’) had high predictive power (AUC range: 0.926–0.958) and were largely parapatric in Madagascar (range overlap *c*. 0.06–0.12; **Fig. 1a**). When combined with the PCA results (**Fig. S2**, **Table S7**) and earlier phytogeographic (Humbert 1955; Dransfield *et al.* 2011) and climate data (Cornet 1974; Yoder & Nowak 2006; Gamisch *et al.* 2016), these niches can be characterized as follows: *A* occupies the seasonally dry Northwest ‘Sambirano’ rainforest (*c.* 0–800 m), maintaining high annual temperature and low annual (but variable seasonal) rainfall; *B* largely coincides with the ‘Eastern Lowlands’ of coastal and humid rainforest (*c.* 0–800 m), featuring high annual rainfall at ‘intermediate’ annual and seasonal temperatures; and niche *C* extends from the eastern slopes to the plateau of the ‘Central Highlands’ (*c.* 800–1,800 m) of sub-humid montane forest and sclerophyllous forest/scrubland, receiving relatively low annual rainfall and low annual (but variable seasonal) temperature.

**Fig. 1.**
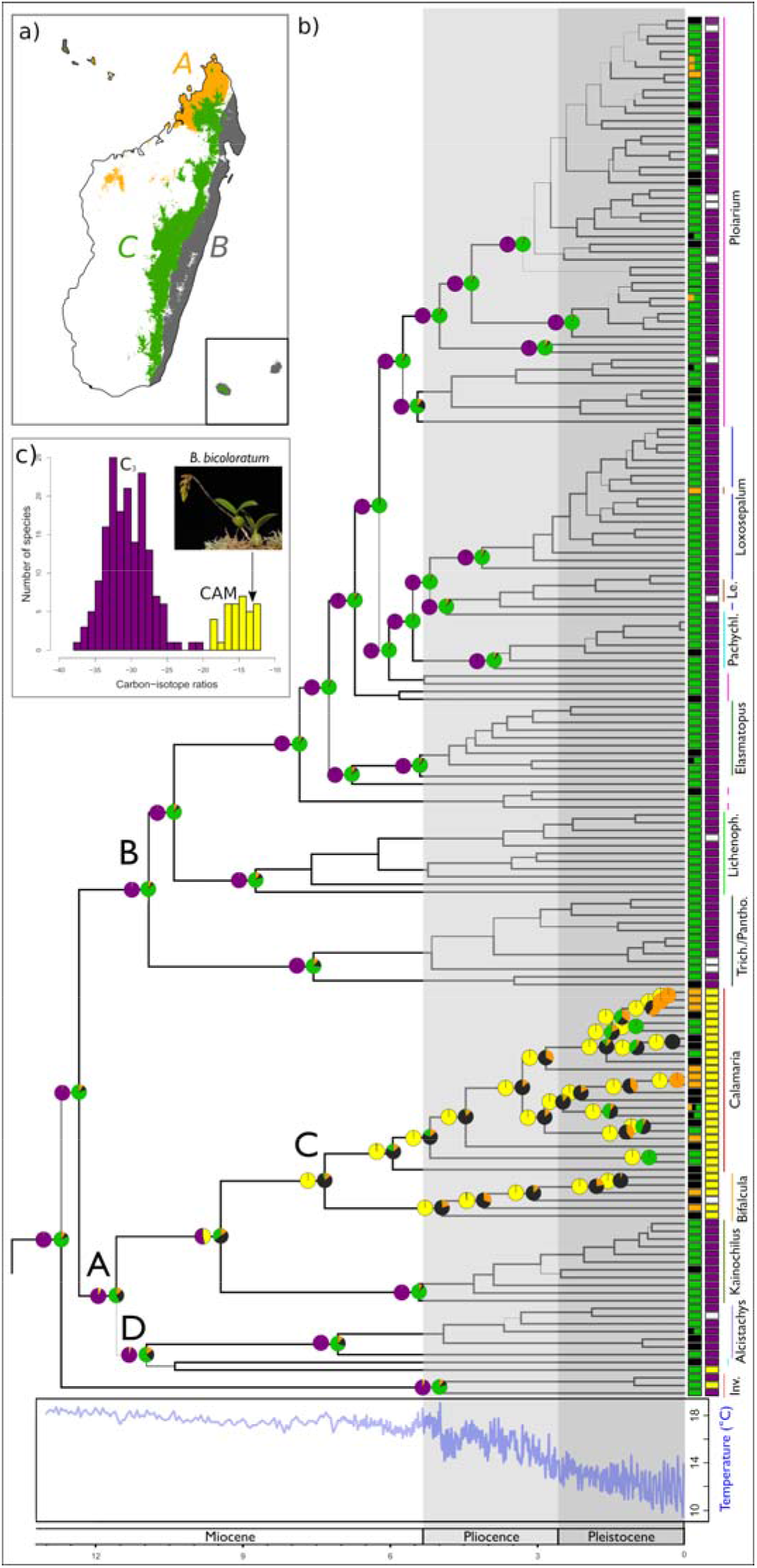
Ancestral climatic niche and photosynthetic pathway reconstructions of Malagasy *Bulbophyllum*. **a)** Current climate condition binary ecological niche models (ENMs) for clusters *A*, *B*, and *C* using the MTSS threshold (inset shows La Réunion and Mauritius). **b**) Ancestral photosynthesis pathway (C_3_, purple; CAM, yellow; no data, white) and niche state reconstructions of Malagasy *Bulbophyllum* (*A*, orange: Northwest ‘Sambirano’; *B*, black: Eastern Lowlands; and *C*, green: Central Highlands), as estimated in Bayestraits and plotted onto the BEAST-derived chronogram (Supplemental **Fig. S1**) with pie charts at selected nodes indicating posterior probability (PP) values. A–D above nodes indicate major clades. **c**) Histogram of frequency of observed carbon-isotope ratios of Malagasy *Bulbophyllum* (see also Supplemental Table **S4**). Inset photograph shows habitus of the CAM species *B. bicoloratum* (sect. *Calamaria*). Photo credit: Rogier van Vugt.

### Ancestral niche states and transition rates

The Bayesian reconstruction of ancestral niche states (*A*, *B*, *C*) along the MCC chronogram (**Fig. 1b**) placed the crown node of Malagasy *Bulbophyllum* (*c*. 12.70 Ma) in the Central Highlands (*C*) with high probability (PP = 81.1), and the same was true for the crown nodes of clades B (PP = 87.2), D (PP = 78.1) and, to a lesser extent, A (PP = 57.5). In fact, most lineages and extant species of clades B and D retained this ancestral niche, with the exception of relatively few and independent transitions on terminal tip branches, mostly from *C* to the Eastern Lowlands (*B*) or, very rarely, from *C* to the Sambirano (*A*). Clade A, however, underwent a notable shift from *C* to *B* during the late Miocene, either coinciding with the origin of subclade C (stem node: PP = 50.6; *c*. 9.45 Ma) or, more likely, with its onset of diversification (crown node: PP = 80.8; *c*. 7.34 Ma); subsequently, relatively frequent transitions occurred (towards the tips) from *B* to *A* or *C* (or very rarely from *A* to *B*), resulting in a subclade with relatively high niche diversity (**Fig. 1b**). Nonetheless, across the entire phylogeny, ancestral niche conservatism (*C*) prevailed, with shifts occurring mainly from *C* to *B* and from *B* to *A* (**Fig. 1b**). In support of this, BAYESTRAITS strongly rejected models opposing these two latter transitions (logBF *q*_CB_ = −6.9; *q*_BA_ = −10.4) but none with zero (uni-directional) transitions (i.e. *q*_BC_, *q*_CA_, *q*_AB_ or *q*_AC_ = 0; logBF = −1.0–0.9; see **Table S4**).

### Reconstruction of ancestral photosynthetic pathways and correlation with niche states

Carbon-isotope ratios of the 201 *Bulbophyllum* accessions surveyed ranged from −37.75 ‰ to −12.09 ‰ (**Table S4**), resulting in a bimodal frequency distribution (**Fig. 1b**). Based on our binary classification (Winter *et al.* 2015), the great majority of the Malagasy species were found to exhibit C_3_ photosynthesis (159/190, *c*. 83.7%), while those engaging in CAM as major mode of carbon acquisition (31/190; *c.* 16.3%) were almost exclusively confined to (and fixed within) subclade C (29/31, *c.* 93.5%; **Fig. 1b; Table S4**). Accordingly, we identified C_3_ as the most likely ancestral state of Malagasy *Bulbophyllum* (crown node: PP_C3_= 98.8), whereas the group’s major shift towards CAM coincided with the diversification of subclade C (crown node: PP_CAM_ = 99.9). Otherwise, CAM evolved independently only twice, namely in *B. petrae* (sect. *Polyradices*; clade A) and *B. cochinealloides* (sect. *Inversiflora*; clade D). Using Bayestraits, we found strong support for correlated evolution (BF = 8.27) of C_3_ and CAM with, respectively, the ancestral Central Highland niche (*C*) and the derived Sambirano/Eastern Lowland niches combined (*AB*).

### Rates of niche evolution vs. lineage diversification through time

For quantifying rates of niche evolution in BAMM, we used the mean PC1 and PC2 scores of each species based on the 3036 locality-climate *dataset 2* (see **Table S6**for component loadings). As for *dataset 1*, the primary loadings on PC1 vs. PC2 captured measures of temperature/precipitation vs. seasonality/isothermality. For the MCC chronogram with or without subclade C, niche evolution through time was consistently higher and more constant for PC1 relative to PC2 (**Fig. 2a, b**). Notably, for the full tree, niche evolution along PC2 only started in the late Miocene (*c.* 8 Ma), almost coincident with the origin of subclade C (*c*. 7.34 Ma), and then steadily increased toward the present. By contrast, in subclade C, niche evolution for PC1 and PC2 was temporally congruent and steadily increased toward the present at higher rates than in the remaining lineage (**Fig. 2c**).

**Fig. 2.**
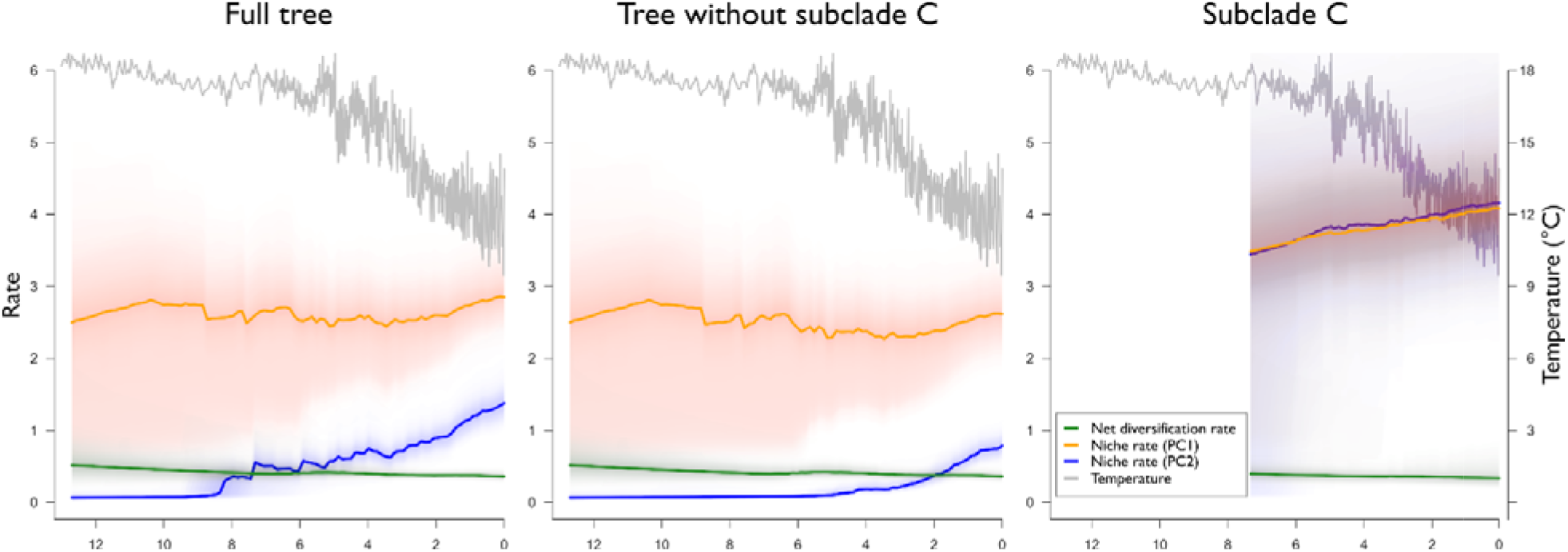
Mean rates of net diversification (*r*) and climatic niche evolution (PC1, PC2) through time. The grey curve indicates global temperature as derived from benthic oxygen isotope ratios (Hansen *et al.* 2013).

Considering lineage diversification, we generally inferred relatively constant, yet slightly declining rates of *r* throughout the history of Malagasy *Bulbophyllum*, regardless of the dataset used (i.e. full tree with or without subclade C, or the latter separately; **Fig. 2a–c**). Moreover, considering HPD intervals, none of the respective diversification parameters (*r*, λ, μ) differed significantly among the three datasets (**Table S7**).

According to RPANDA/DDD, a time-dependent, pure birth model of diversification, with constant speciation and no extinction, provided the best fit for each of these three datasets (**Table S5**). We note that models of environmental-dependent (palaeo-temperature) and diversity-dependent (DD) speciation had essentially equivalent fit (< 2 ΔAICc units) for the full tree with or without subclade C included (**Table S5**). In both instances, however, the estimated carrying capacity (*K*) was almost two to three times the current species richness (*K* = 555/210 and 400/171 spp., respectively). Overall, these results indicate mostly constant speciation with zero extinction throughout the group’s history, even though minor effects of temperature and/or diversity on diversification cannot be fully excluded.

According to STRAPP, there was no significant relationship between net diversification rate (*r*) and niche state (*A*, *B*, *C*) or multi-dimensional climate (PC1, PC2) space, whether estimated for the entire tree or subclade C (all *P* = 0.902–0.998). In sum, these results indicate that the group diversified almost independently of a particular (qualitative or quantitative) niche state (see also **Fig. 2**).

## Discussion

### Spatial-temporal history of Malagasy *Bulbophyllum*

Based on a multi-locus plastid/nuclear phylogeny, we dated the crown age of Malagasy *Bulbophyllum* to the mid-Miocene, *c*. 12.70 Ma (95% HPD: 10.39–15.27 Ma) (**Fig. 1**). This timing is very similar to our previous estimate inferred from a single nuclear (nrITS) marker [*c.* 12.36 (8.23–18.84) Ma; Gamisch & Comes 2019]. Hence, the present results support our previous hypothesis that *Bulbophyllum* colonized Madagascar following the establishment of the Indian Summer monsoon (*c*. 12.9–7.0 Ma) via a single long-distance dispersal event from the Asian-Pacific region (Buerki *et al.* 2013; Gupta *et al.* 2015; Gamisch & Comes 2019, and references therein). Concomitantly, our ancestral niche reconstructions (**Fig. 1**) show that the radiation of *Bulbophyllum* in Madagascar began in interior sub-humid environments of the ‘Central Highlands’ (niche *C*), providing suitable and ‘empty’ (e.g. non-competitive) ecological space required for successful island colonization (Gillespie 2007). Since colonizers likely share similar niche conditions with their source populations (Wiens & Graham 2005; Wiens *et al.* 2010; Steinbauer *et al.* 2016), the ancestor of Malagasy *Bulbophyllum* probably originated from similar habitats in the Asian-Pacific region (e.g. mountains of the Sunda Islands, New Guinea Highlands, India, Indochina and South China). In turn, these results dismiss the hypothesis that insular radiations typically begin in marginal (coastal) habitats fringing the island (Liebherr & Hajek 1990; Schluter 2000), as recently shown for Malagasy *Canarium* (Burseraceae; Federman *et al.* 2015).

### Major transition from C3 to CAM, associated with invasion of high rainfall forest

Our results clearly identify C_3_ as the ancestral and predominant mode of photosynthesis in Malagasy *Bulbophyllum* (159/190 spp., *c*. 83.7%), while strong CAM is relatively rare (31/190,; *c.* 16.3%) and mostly confined to subclade C (29/31, *c.* 93.5%), except for two independent, species-specific transitions (*B. petrae*, *B. cochinealloides*; **Fig. 1b**, **Table S4**). Thus, the majority of species still occupying the ancestral ‘Central Highlands’ niche (*C*) exhibit C_3_ photosynthesis, while CAM is mostly associated with relatively rare niche shifts to either the ‘Eastern Lowlands’ (*B*) or the ‘Sambirano’ (*A*), as mostly found in subclade C (**Fig. 1b**). Hence, Malagasy *Bulbophyllum* features considerable ecological niche (*C*) and physiological (C_3_) conservatism, yet these phylogenetic constraints (*sensu* Wiens *et al.* 2010) were apparently not strong enough to restrain the correlated evolution of alternative (derived) niche states (*A*, *B*) and CAM. More specifically, the group’s major transition from C_3_ to CAM most likely occurred at the crown node of subclade C (PP = 99.9), i.e. during the late Miocene (*c*. 7.36 Ma; 95% HPD: 5.83–8.89 Ma), and was accompanied by a major niche shift from the sub-humid Central Highlands (*C*) to the warmer and moister Eastern Lowlands (*B*), followed by colonizations of the hotter/seasonally dry ‘Sambirano’ (*A*) or reversals back into the highlands (**Fig. 1b, Table S3**).

Based on these ancestral niche reconstructions, it appears unlikely that CAM initially evolved in Malagasy *Bulbophyllum* as an adaptive response to dry environments or epiphytism *per se* (e.g. Gravendeel *et al.* 2004; Silvera *et al.* 2009, 2010; Bone *et al.* 2015). Instead, these data suggest that CAM initially served as a ‘gateway’ trait (*sensu* Donoghue & Edwards 2014; Bouchenak-Khelladi *et al.* 2015), facilitating the invasion of a novel adaptive zone of high rainfall coastal forest (*B*). Here, CAM could have been selectively advantageous to compensate for the wetting of leaves (stomata blockage) while absorbing more CO_2_ when subsequently dry and/or over a longer time than the photoperiod (Pierce *et al*. 2002; Lüttge 2004; see also Keeley 1998 for similar explanations of CAM in aquatic plants). Subsequently, CAM likely pre-adapted species of those wetter (*B*) habitats to the seasonally dry ‘Sambirano’ rainforest (*A*), further widening the spatial-ecological amplitude of Malagasy *Bulbophyllum*.

But what extrinsic (e.g. climatic) factors could have caused, or at least promoted these niche and physiological transitions in subclade C? Rapid aridification under low CO_2_ conditions in the mid-to-late Miocene is frequently cited as potential factor promoting the evolution of CAM (e.g. Bone *et al.* 2015; Hu 2018; Li *et al.* 2019). In fact, this timing broadly matches the late Miocene diversification of subclade C and its corresponding shifts towards CAM and into the Eastern Lowlands (see above). However, this timing also matches a period when the Indian Summer monsoon, whose seasonal precipitation influx is thought to have created the Northwest ‘Sambirano’ from *c*. 8 Ma onwards (Yoder & Nowak 2006; Buerki *et al.* 2013), reached its full strength (*c*. 7 Ma; Gupta *et al.* 2015). Hence, these changes in monsoon climate could have further created environmental instability across the comparatively older sub-humid and humid forest biomes of the Central Highlands and Eastern Coast of Madagascar, respectively (Cornet 1974; Buerki *et al.* 2013). In turn, these climatic vicissitudes during the late Miocene might have especially favoured *Bulbophyllum* species with “highly flexible CAM” (cf. Pierce *et al.* 2002) to move from sub-humid highland to wetter or seasonally dry lowland niches.

### Rates of species diversification and niche evolution in relation to CAM

According to adaptive radiation theory (Simpson 1944; Schluter 2000), linages exposed to ecological opportunity, and especially after island colonizations (Gillespie 2007), are expected to show an ‘early-burst’ of both species diversification and phenotypic evolution, followed by a slowdown in these rates due to limits imposed by geographical area, species competition and/or niche availability (Harmon *et al.* 2008; Rabosky 2009; Silvestro *et al.* 2013; Givnish *et al.* 2014; Herrera 2017; Linder & Bouchenak-Khelladi 2017; Burbrink *et al.* 2019). In Madagascar, such patterns of declining diversification rates have been reported in *Angraecum* orchids (Andriananjamanantsoa *et al.* 2016) and several amphibian/reptile and bird lineages (e.g. Jønsson *et al.* 2012; Reddy *et al.* 2012; Scantlebury 2013).

However, our rate-through-time analyses for Malagasy *Bulbophyllum* (**Fig. 2**) indicate (1) mostly constant rates of speciation with little or no extinction in Malagasy *Bulbophyllum*(with full tree mean μ = 0.06), including no evidence of strong density dependence (**Table S5**); and (2) elevated rates of niche (PC2) evolution within the last *c*. 8 million years, largely reflecting the widened climatic niche space of the CAM-possessing subclade C (in terms of seasonality/isothermality; **Table S6**). Together, these results indicate that limits for both species diversity and niche evolution have not yet been reached in this hyper-diverse radiation (Rabosky 2009), despite its relative antiquity (late Miocene) and insular setting. We suspect this lack of spatial-ecological boundaries in this group could be caused by a combination of factors, including general features of tropical orchids (e.g. low population sizes: McCormick & Jacquemyn 2014; Hu *et al.* 2017; vast surface area of their mostly epiphytic life zone: Gravendeel *et al.* 2004), lineage-specific characteristics (e.g. relative unspecialized pollination by a single functional group of fly pollinators: Fischer *et al*. 2007; Gamisch *et al.* 2014) and, perhaps foremost, historical contingencies (e.g. Moen & Morlon 2014), especially the evolution of CAM, providing a mechanism to escape from the ancestrally highland niche (*C*) into a wider range of lowland (*A*, *B*) niches (see above).

However, despite these continuing ecological opportunities, we found no significant differences in diversification parameter estimates (λ, *μ*, *r*) between the CAM-possessing subclade C and the full tree with or without this lineage (**Table S7**). Such lack of association between CAM and diversification has also been reported in Afro-Malagasy Eulophiinae (Bone *et al.* 2015) and Asian *Bulbophyllum* (Hu 2018), thereby challenging the idea of CAM as a ‘key innovation’ in tropical orchids (e.g. Gravendeel *et al*. 2004; Silvestro *et al*. 2013; Bouchenak-Khelladi *et al*. 2015; Givnish *et al.* 2014, 2015). However, we are not aware of similar studies reporting not only near constant diversification rates in a large endemic Malagasy plant lineage, but also a major CAM-mediated niche shift from comparatively drier to moister habitats (here form niches *C* to *B*), whether in this island or elsewhere.

## Conclusions

The likely single species of *Bulbophyllum* colonizing Madagascar in the mid-Miocene (12.70 Ma) arrived in the Central Highland region and exhibited C_3_ photosynthesis. These niche and metabolic states are retained by the great majority of extant species. Nonetheless, a major shift from C_3_ to CAM, coinciding with the crown age of subclade C (*c*. 7.36 Ma), provided a mechanism to escape from the ancestrally conserved sub-humid Central Highland niche, and to first move into the Eastern Lowland rainforests, and then into the seasonal dry Northwest ‘Sambirano’. Hence, CAM in these mostly epiphytic orchids is best characterised as a ‘gateway’ trait, first conveying a selective advantage in coastal forest habitats with high rainfall (e.g. possibly compensating for wetted leaves/blocked stomata), and then pre-adapting such lowland species to further colonize seasonally dry habitats. Although the shift towards CAM resulted in elevated rates of climatic niche evolution, it had no effect on species diversification. Nevertheless, evolutionary change in this single physiological trait had a major role in shaping the evolution of Madagascar’s most species-rich orchid genus by increasing its overall spatial-ecological amplitude and ‘total diversification’ viz. species richness (see also Rabosky 2017).

## Acknowledgements

We are grateful to the members of the Botanical Garden of Vienna University (Michael Kiehn, Anton Sieder, David Prehsler) and the Parc Botanique et Zoologique de Tsimbazaza (Solo Rapanarivo, Jacky Andriantiana) for general support and help with fieldwork in Madagascar; the Département des Eaux et Fôrets (Madagascar) for issuing collecting permits; the curators of herbaria (G, K, MO, NEU, P, SZU, TAN, W, WU) for the loan of specimens; Johan Hermans and Claire Hermans (Royal Botanical Gardens, Kew, UK) for help in species identification; Myriam Gaudeul and the Muséum national d'Histoire naturelle (Paris, France), Tariq Stévart (Missouri Botanical Garden, St. Louis, USA) and Simon Verlynde (The Graduate Center, CUNY, USA) for providing leaf samples for DNA extraction; and to Benjamin L. Turner and Dayana Agudo (Smithsonian Tropical Research Institute, Panama City, Panama) for providing facility and aid in running the IRMS analyses. This study was funded by the FWF (Austrian Science Fund) grant P29371 to Hans Peter Comes.

## Author Contributions

A.G. and H.P.C. conceived and designed the study. A.G. performed the analyses. A.G analysed the data. K.W. supervised the IRMS analyses. G.F. contributed sequences. A.G., H.P.C. and K.W. interpreted the data. A.G. and H.P.C drafted the manuscript. All authors read and approved the final manuscript.

## Competing Interests statement

The authors declare no competing interests.

## Notes

### Competing Interest Statement

The authors have declared no competing interest.

