## Supplemental for "Evolution of Crassulacean Acid Metabolism (CAM) as an escape from ecological niche conservatism in Malagasy *Bulbophyllum* (Orchidaceae)"

**Supplement**

**Including Supplemental Tables S1–S7, Figs. S1–S2**

**Supplemental Tables**

**Table S1. Species and markers used in the phylogenetic study of Malagasy *Bulbophyllum*.** GenBank accessions numbers are provided for all published sequences (newly added sequences are marked in bold). Recently described species (sensu Hermans *et al*., in press) are indicated by asterisks.

| **Species** | ***Section*** | **Collection** | ***atp*H** | **nrITS** | ***psb*A** | ***trn*D** | ***ycf*1** | ***Mat*k** | ***PI*** | ***Xdh*** |
| --- | --- | --- | --- | --- | --- | --- | --- | --- | --- | --- |
| *B. bathieanum* | *Alcistachys* | FS5219 | **MN618476** | MH822368 | **MN618894** | **MN618946** | **MN619145** | **MN618638** | **MN618777** | **MN619020** |
| *B. brevipetalum* | *Alcistachys* | FS2048 | - | - | - | - | **MN619146** | **MN618641** | - | **MN619021** |
| *B. hamelinii* | *Alcistachys* | FS1085 | **MN618496** | EF195979.1 | EF200380.1 | EF202195.1 | **MN619164** | **MN618662** | **MN618801** | **MN619041** |
| *B. occlusum* | *Alcistachys* | FS722 | **MN618533** | EF196040.1 | EF200381.1 | EF202196.1 | **MN619197** | **MN639625** | **MN618838** | **MN619080** |
| *B. sp. nov1* | *Alcistachys* | FS4310 | **MN618560** | **MN618610** | **MN618926** | **MN618991** | **MN619224** | **MN618737** | **MN618867** | **MN619108** |
| *B. sulfureum* | *Alcistachys* | FS1585 | **MN618570** | EF196071.1 | EF200382.1 | EF202197.1 | **MN619235** | **MN618751** | **MN618878** | **MN619121** |
| *B. variegatum* | *Alcistachys* | FS799 | KJ558987 | KJ558713.1 | EF200379.1 | EF202194.1 | KJ558557.1 | **MN618758** | KJ558765 | **MN639650** |
| *B. capuronii* | *Bifalcula* | FS1010 | KJ558989 | EF195966.1 | EF200383.1 | KJ558661.1 | KJ558559.1 | **MN639613** | KJ558773 | **MN619022** |
| *B. complanatum* | *Bifalcula* | FS1298 | KJ558964 | EF633602.1 | KJ558910.1 | KJ558667.1 | KJ558534.1 | **MN639614** | KJ558785 | **MN619028** |
| *B. implexum* | *Bifalcula* | FS1207 | KJ558960 | EF196031.1 | EF200385.1 | EF202199.1 | KJ558531.1 | **MN618673** | KJ558781 | **MN619049** |
| *B. minutum* | *Bifalcula* | FS1006 | KJ558986 | EF196061.1 | EF200386.1 | EF202198.1 | KJ558556.1 | **MN639622** | KJ558771 | **MN619071** |
| *B. sp. nov1* | *Bifalcula* | FS2707 | KJ558968 | KJ558722.1 | KJ558912.1 | KJ558669.1 | KJ558538.1 | **MN618738** | KJ558789 | **MN619109** |
| *B. sp. nov3* | *Bifalcula* | FS5534 | KJ558970 | KJ558740.1 | KJ558930.1 | KJ558687.1 | KJ558540.1 | **MN618742** | KJ558807 | **MN619112** |
| *B. ambongense* | *Calamaria* | FS6023 | KJ559006 | KJ558755.1 | KJ558946.1 | KJ558703.1 | KJ558576.1 | **MN618626** | KJ558823 | **MN619009** |
| *B. cirrhoglossum* | *Calamaria* | FS4125 | KJ558971 | KJ558728.1 | KJ558918.1 | **MN618952** | KJ558541.1 | **MN618647** | KJ558795 | **MN619025** |
| *B. elliotii* | *Calamaria* | FS4137 | KJ558972 | KJ558729.1 | KJ558919.1 | KJ558676.1 | KJ558542.1 | - | KJ558796 | **MN619038** |
| *B. erectum* | *Calamaria* | FS1022 | KJ558985 | KJ558716.1 | KJ558905.1 | KJ558662.1 | KJ558555.1 | **MN618657** | KJ558774 | **MN619039** |
| *B. hildebrandtii* | *Calamaria* | FS1169 | KJ558958 | KJ558720.1 | KJ558909.1 | KJ558666.1 | KJ558529.1 | - | KJ558779 | **MN619043** |
| *B. humblotii* | *Calamaria* | FS1008 | KJ558973 | EF195986.1 | EF200402.1 | EF202216.1 | KJ558543.1 | **MN618670** | KJ558772 | **MN619047** |
| *B. incurvum* | *Calamaria* | REU2 | KJ559021 | KJ558761.1 | KJ558952.1 | KJ558709.1 | KJ558591.1 | **MN618674** | KJ558830 | **MN619050** |
| *B. lecouflei* | *Calamaria* | FS2715_FS1278 | KJ558963 | KJ558723.1 | KJ558913.1 | KJ558670.1 | KJ558539.1 | - | KJ558784 | **MN619059** |
| *B. luteobracteatum* | *Calamaria* | FS5512_OR1257 | KJ559004 | KJ558737.1 | KJ558927.1 | KJ558684.1 | KJ558574.1 | **MN618688** | KJ558804 | **MN619065** |
| *B. maculatum cf* | *Calamaria* | FS1133 | KJ558956 | EF195983.1 | EF200392.1 | EF202206.1 | KJ558527.1 | **MN618690** | KJ558777 | **MN619068** |
| *B. malawiense* | *Calamaria* | OR146_05 | KJ558984 | KJ558759.1 | KJ558950.1 | KJ558707.1 | KJ558554.1 | **MN618691** | KJ558828 | **MN619069** |
| *B. obtusatum* | *Calamaria* | FS1170 | KJ558959 | EF196039.1 | EF200395.1 | EF202209.1 | KJ558530.1 | **MN639624** | KJ558780 | **MN619079** |
| *B. occultum* | *Calamaria* | FS603 | KJ558990 | KJ558760.1 | KJ558951.1 | KJ558708.1 | KJ558560.1 | - | KJ558829 | **MN619081** |
| *B. pentastachium* | *Calamaria* | FS827 | KJ558983 | KJ558715.1 | KJ558904.1 | KJ558660.1 | KJ558553.1 | **MN618714** | KJ558769 | **MN619089** |
| *B. pervillei* | *Calamaria* | FS818 | KJ558981 | EF196049.1 | EF200400.1 | EF202214.1 | KJ558551.1 | **MN618715** | KJ558767 | **MN619090** |
| *B. pusillum* | *Calamaria* | FS1459 | KJ558965 | EF196057.1 | EF200398.1 | EF202212.1 | KJ558535.1 | **MN618722** | KJ558786 | **MN619096** |
| *B. rubrum* | *Calamaria* | FS5603_OR1259 | KJ559005 | KJ558741.1 | KJ558931.1 | KJ558688.1 | KJ558575.1 | - | KJ558808 | **MN619100** |
| *B. ruginosum* | *Calamaria* | FS5885_OR1400 | KJ559017 | KJ558751.1 | KJ558941.1 | KJ558698.1 | KJ558587.1 | **MN618724** | KJ558818 | **MN619101** |
| *B. sambiranense* var. *latibracteatum* | *Calamaria* | FS5719 | KJ559015 | KJ558746.1 | KJ558936.1 | KJ558693.1 | KJ558585.1 | **MN618725** | KJ558813 | **MN619102** |
| *B. sp. nov1* | *Calamaria* | FS5246 | KJ559011 | KJ558731.1 | KJ558921.1 | KJ558678.1 | KJ558581.1 | **MN618739** | KJ558798 | **MN619110** |
| *B. trifarium* | *Calamaria* | FS1224 | KJ558961 | EF196072.1 | EF200399.1 | EF202213.1 | KJ558532.1 | **MN618754** | KJ558782 | **MN619124** |
| *B. bicoloratum* | *Calamaria* | FS1052 | KJ558980 | EF195964.1 | EF200387.1 | EF202200.1 | KJ558550.1 | **MN618639** | KJ558827 | **MN639634** |
| *B. histrionicum* | *Calamaria* | FS1232 | KJ558962 | EF196062.1 | EF200397.1 | EF202211.1 | KJ558533.1 | **MN618666** | KJ558783 | **MN639638** |
| *B. senghasii* | *Calamaria* | FS3969 | KJ558976 | KJ558726.1 | KJ558916.1 | KJ558673.1 | KJ558546.1 | - | KJ558793 | **MN639646** |
| *B. francoisii* | *Elasmatopus* | FS691 | - | EF195977.1 | EF200411.1 | EF202221.1 | **MN619162** | **MN618660** | - | - |
| *B. amphorimorphum* | *Elasmatopus* | K16926 | **MN618467** | EF196069.1 | EF200418.1 | EF202228.1 | **MN619137** | **MN618630** | **MN618768** | **MN619012** |
| *B. analamazoatrae* | *Elasmatopus* | FS773 | **MN618468** | EF195956.1 | EF200408.1 | EF202217.1 | **MN619138** | **MN618631** | **MN618769** | **MN619013** |
| *B. aubrevillei* | *Elasmatopus* | FS620 | **MN618474** | EF195961.1 | **MN618892** | EF202219.1 | **MN619142** | **MN618635** | **MN618775** | **MN619018** |
| *B. latipetalum* | *Elasmatopus* | FS688 | **MN618509** | **MN618592** | EF200415.1 | EF202225.1 | **MN619177** | **MN618681** | **MN618814** | **MN619057** |
| *B. oxycalyx* | *Elasmatopus* | FS1073 | **MN618539** | EF196044.1 | EF200416.1 | EF202226.1 | **MN619202** | **MN618710** | **MN618843** | **MN619086** |
| *B. pandurella* | *Elasmatopus* | S6641 | **MN618541** | **MN618601** | **MN618919** | **MN618979** | **MN619204** | **MN618712** | **MN618845** | **MN619088** |
| *B. rauhii* | *Elasmatopus* | FS1326 | **MN618551** | EF196054.1 | EF200413.1 | EF202223.1 | **MN639665** | **MN639629** | **MN618857** | **MN619098** |
| *B. sp.* | *Elasmatopus* | FS4071 | **MN618565** | **MN618604** | **MN618925** | **MN618990** | - | **MN618736** | **MN618866** | **MN619107** |
| *B. sp. nov9* | *Elasmatopus* | FS5430 | **MN618563** | **MN618614** | **MN618928** | **MN618994** | **MN619227** | **MN618744** | **MN618871** | **MN619114** |
| *B. cardiobulbum* | *Inversiflora* | FS1641 | KJ558966 | KJ558721.1 | KJ558911.1 | KJ558668.1 | KJ558536.1 | **MN618645** | KJ558787 | **MN619023** |
| *B. cochinealloides** | *Inversiflora* | JH6718 | **MN618483** | **MN618584** | **MN618899** | **MN618954** | **MN619151** | **MN618649** | **MN618787** | **MN619027** |
| *B. uroplatoides** | *Inversiflora* | FS5055 | **MN618574** | **MN618619** | **MN618935** | **MN619002** | **MN619240** | **MN618757** | **MN618881** | **MN639649** |
| *B. mirificum* | *Kainochilus* | LG4797 | - | **MN618596** | - | **MN618972** | **MN648670** | **MN639623** | **MN618827** | - |
| *B. alexandreae* | *Kainochilus* | FS779 | **MN618463** | EF195952.1 | EF200419.1 | EF202229.1 | **MN619132** | **MN618624** | **MN618764** | **MN619007** |
| *B. anjozorobeense* | *Kainochilus* | FS1457 | **MN618470** | EF195958.1 | EF200420.1 | EF202230.1 | **MN619139** | **MN618632** | **MN618771** | **MN619015** |
| *B. cylindrocarpum* | *Kainochilus* | REU7 | **MN618489** | **MN618585** | **MN618901** | **MN618956** | **MN619154** | **MN618652** | **MN618792** | **MN619033** |
| *B. edentatum* | *Kainochilus* | FS866 | - | EF195972.1 | EF200421.1 | EF202231.1 | **MN619160** | - | **MN618798** | **MN619037** |
| *B. horizontale* | *Kainochilus* | AJL148 | KJ558955 | KJ558712.1 | EF200422.1 | EF202232.1 | KJ558526.1 | **MN618667** | KJ558764 | **MN619045** |
| *B. imerinense* | *Kainochilus* | FS901 | **MN618500** | EF195987.1 | EF200423.1 | EF202233.1 | **MN619170** | **MN618672** | **MN618805** | **MN619048** |
| *B. jeanbosseri** | *Kainochilus* | ## | **MN618504** | **MN618591** | **MN618906** | **MN618963** | **MN639659** | **MN618676** | **MN618809** | **MN619053** |
| *B. multiligulatum* | *Kainochilus* | FS5612 | **MN639606** | **MN618597** | **MN618914** | **MN618973** | **MN619190** | **MN618700** | **MN618831** | **MN619075** |
| *B. sp. nov1* | *Kainochilus* | FS5554 | **MN618561** | **MN618609** | - | - | **MN619225** | **MN618740** | **MN618868** | **MN619111** |
| *B. vigueri* | *Kainochilus* | FS2580 | **MN618578** | **MN618620** | **MN618937** | **MN619004** | **MN619244** | **MN618761** | **MN618885** | **MN619129** |
| *B. jumelleanum* | *Lepiophylax* | FS5791 | **MN618505** | MH822391 | **MN639611** | **MN618964** | **MN619173** | **MN618677** | **MN618810** | **MN619054** |
| *B. sp1* | *Lepiophylax* | FS3197 | **MN618555** | **MN618606** | - | **MN618986** | - | **MN618730** | **MN618862** | **MN619104** |
| *B. trilineatum* | *Lepiophylax* | FS4776 | **MN618573** | **MN618618** | **MN618934** | **MN619001** | **MN619238** | **MN618755** | **MN618880** | **MN619125** |
| *B. calyptropus* | *Lepiophylax* | FS2202 | **MN618480** | EF195965.1 | EF200427.1 | EF202238.1 | **MN619149** | **MN618644** | **MN618783** | **MN639636** |
| *B. forsythianum* | *Lichenophylax* | FS1552 | - | EF196063.1 | EF200425.1 | EF202236.1 | **MN639655** | **MN618659** | - | - |
| *B. mangenotii* | *Lichenophylax* | FS1608 | - | MH822398 | - | **MN618970** | **MN619184** | **MN618692** | - | - |
| *B. amparihibense* | *Lichenophylax* | FS4365 | - | MH822367 | **MN618889** | **MN618941** | **MN619136** | **MN618629** | **MN618767** | - |
| *B. bryophytoides* | *Lichenophylax* | FS2595 | **MN618478** | MH822371 | **MN618896** | **MN618949** | **MN619147** | **MN618642** | **MN618780** | - |
| *B. debile* | *Lichenophylax* | FS4070 | **MN618490** | MH822381 | **MN639610** | **MN618957** | **MN619155** | **MN639617** | **MN618793** | - |
| *B. lakatoense* | *Lichenophylax* | FS1017 | **MN618507** | MH822393 | **MN618908** | **MN618965** | **MN619175** | **MN618679** | **MN618812** | - |
| *B. lichenophylax* | *Lichenophylax* | FS5529 | **MN618512** | **MN618593** | - | - | **MN639660** | - | **MN618818** | - |
| *B. neglectum* | *Lichenophylax* | MO3984 | **MN618528** | **MN618598** | - | - | **MN619193** | - | **MN618835** | - |
| *B. quinquecornutum* | *Lichenophylax* | FS709 | **MN639608** | EF196060.1 | - | EF202235.1 | - | - | **MN618856** | - |
| *B. afzelii* | *Lichenophylax* | FS2526 | **MN618461** | MH822365 | **MN618887** | **MN618939** | **MN619130** | **MN618622** | **MN639668** | - |
| *B. hapalanthos* | *Lichenophylax* | FS880 | **MN639605** | EF195980.1 | EF200424.1 | EF202234.1 | **MN639656** | - | **MN639671** | - |
| *B. sciaphile* | *Loxosepalum* | FS769 | - | EF196059.1 | EF200438.1 | **MN618985** | **MN639666** | **MN618728** | - | - |
| *B. florulentum* | *Loxosepalum* | FS650 | - | EF195996.1 | EF200444.1 | EF202254.1 | **MN619161** | **MN618658** | **MN618799** | - |
| *B. alleizettei* | *Loxosepalum* | FS746 | **MN618464** | EF195953.1 | EF200458.1 | EF202268.1 | **MN619133** | **MN618625** | **MN618765** | **MN619008** |
| *B. ambrense* | *Loxosepalum* | FS5886 | **MN618465** | **MN618579** | **MN618888** | **MN618940** | **MN619134** | **MN618627** | **MN639669** | **MN619010** |
| *B. amoenum* | *Loxosepalum* | FS932 | **MN618466** | EF195955.1 | EF200426.1 | EF202237.1 | **MN619135** | **MN618628** | **MN618766** | **MN619011** |
| *B. approximatum cf* | *Loxosepalum* | FS4328 | **MN618473** | **MN618581** | - | **MN618944** | **MN619141** | **MN639612** | **MN618774** | **MN619017** |
| *B. baronii* | *Loxosepalum* | OR56 | **MN618475** | **MN618582** | **MN618893** | **MN618945** | **MN619144** | **MN618637** | **MN618776** | **MN619019** |
| *B. conchidioides* | *Loxosepalum* | FS682 | **MN618485** | EF195969.1 | EF200428.1 | EF202239.1 | **MN648669** | **MN639615** | **MN618789** | **MN619029** |
| *B. decaryanum* | *Loxosepalum* | FS949 | **MN618491** | EF195946.1 | EF200441.1 | EF202251.1 | **MN619156** | **MN639618** | **MN618794** | **MN619034** |
| *B. densum* | *Loxosepalum* | REU4 | **MN618492** | MH822382 | **MN618902** | - | **MN619157** | **MN618653** | **MN618795** | **MN619035** |
| *B. leandrianum* | *Loxosepalum* | FS1028 | **MN618510** | EF196021.1 | EF200429.1 | EF202240.1 | - | **MN618682** | **MN618815** | **MN619058** |
| *B. leptostachyum* | *Loxosepalum* | FS874 | **MN618511** | EF196005.1 | EF200450.1 | EF202260.1 | **MN619179** | **MN639620** | **MN618817** | **MN619060** |
| *B. lineariligulatum* | *Loxosepalum* | FS1370 | **MN618513** | **MN618594** | **MN618910** | **MN618968** | **MN639661** | **MN618684** | **MN618819** | **MN619061** |
| *B. multiflorum* | *Loxosepalum* | FS1079 | **MN618524** | EF196027.1 | EF200431.1 | EF202243.1 | **MN619189** | **MN618699** | **MN639672** | **MN619074** |
| *B. nutans* | *Loxosepalum* | FS971 | **MN618531** | EF196036.1 | EF200434.1 | EF202245.1 | **MN619196** | **MN618706** | **MN639673** | **MN619078** |
| *B. sphaerobulbum* | *Loxosepalum* | 128A479 | **MN618566** | - | **MN618930** | **MN618997** | **MN619232** | **MN618748** | **MN618875** | **MN619118** |
| *B. vakonae* | *Loxosepalum* | FS871 | **MN618575** | EF196042.1 | EF200437.1 | EF202248.1 | **MN619241** | - | **MN618882** | **MN619126** |
| *B. variifolium* | *Loxosepalum* | FS2532 | **MN618576** | MH822429 | **MN618936** | **MN619003** | **MN619242** | **MN618759** | **MN618883** | **MN619127** |
| *B. melleum* | *Loxosepalum* | JH5483 | **MN618520** | EF196030.1 | EF200430.1 | EF202241.1 | **MN619186** | **MN618695** | **MN618826** | **MN639641** |
| *B. sp16* | *Loxosepalum* | FS683 | **MN618559** | - | EF200442.1 | EF202252.1 | **MN619222** | **MN618735** | **MN618865** | **MN639648** |
| *B. lyperocephalum* | *Lyperocephalum* | FS2680 | **MN618516** | MH822396 | - | **MN618969** | **MN619182** | **MN618689** | **MN618822** | **MN619066** |
| *B. ikongoense* | *Lyperostachys* | FS1514 | **MN618499** | **MN618589** | EF200461.1 | EF202271.1 | **MN619169** | **MN618671** | **MN618804** | - |
| *B. moratii* | *Pachychlamys* | FS4250, FS4297 | **MN618523** | MH822403 | **MN618913** | **MN635570** | **MN639664** | **MN618698** | **MN618830** | - |
| *B. liparidioides* | *Pachychlamys* | FS706 | - | EF196022.1 | EF200462.1 | EF202272.1 | **MN639662** | **MN618685** | - | **MN619062** |
| *B. longivaginans* | *Pachychlamys* | FS1402 | **MN618514** | EF196025.1 | EF200463.1 | EF202273.1 | **MN619180** | **MN618686** | **MN618820** | **MN619063** |
| *B. macrocarpum* | *Pachychlamys* | REU5 | **MN618517** | **MN618595** | **MN618911** | - | **MN619183** | **MN639621** | **MN618823** | **MN619067** |
| *B. molossus* | *Pachychlamys* | FS622 | **MN618521** | EF196032.1 | EF200465.1 | EF202275.1 | **MN619187** | **MN618696** | **MN618828** | **MN619072** |
| *B. pachypus* | *Pachychlamys* | FS986 | **MN618540** | EF196047.1 | EF200466.1 | EF202276.1 | **MN619203** | **MN618711** | **MN618844** | **MN619087** |
| *B. sandrangatense* | *Pachychlamys* | FS661 | **MN618553** | EF196058.1 | EF200467.1 | EF202277.1 | **MN619214** | **MN618726** | **MN618860** | **MN619103** |
| *B. sp. nov3* | *Pachychlamys* | FS2562 | **MN618562** | **MN618612** | **MN618927** | **MN618993** | **MN619226** | **MN618743** | **MN618870** | **MN619113** |
| *B. vestitum* | *Pachychlamys* | FS631 | **MN618577** | EF196076.1 | EF200469.1 | EF202279.1 | **MN619243** | **MN618760** | **MN618884** | **MN619128** |
| *B. bryophilum* | *Pantoblepharon* | FS1409 | - | - | - | **MN618948** | - | - | **MN618779** | - |
| *B. discilabium* | *Pantoblepharon* | 279A476 | **MN618493** | **MN618586** | **MN618903** | **MN618958** | **MN619158** | **MN618654** | **MN618796** | - |
| *B. duplicatum** | *Ploiarium* | P00097314 | **MN618532** | - | - | **MN618960** | - | - | - | - |
| *B. paleiferum* | *Ploiarium* | FS3254 | - | MH822408 | - | **MN618978** | - | - | - | - |
| *B. abbreviatum* | *Ploiarium* | MO4836204 | - | - | - | **MN618938** | **MN639651** | **MN618621** | - | - |
| *B. auriflorum* | *Ploiarium* | K16655 | - | EF195963.1 | EF200473.1 | EF202283.1 | **MN619143** | **MN618636** | - | - |
| *B. humbertii* | *Ploiarium* | K16929 | - | EF195985.1 | EF200477.1 | EF202287.1 | **MN619168** | **MN618669** | - | - |
| *B. leptochlamys* | *Ploiarium* | K16624 | - | EF195971.1 | EF200479.1 | EF202289.1 | **MN619178** | **MN618683** | - | - |
| *B. septatum cf* | *Ploiarium* | FS3151 | - | **MN618603** | - | **MN635571** | **MN619216** | **MN618729** | - | - |
| *B. sp9* | *Ploiarium* | FS1558 | - | EF195991.1 | EF200501.1 | EF202313.1 | **MN619219** | **MN618732** | - | - |
| *B. sp15* | *Ploiarium* | FS602 | - | EF196066.1 | - | EF202307.1 | **MN619221** | **MN618734** | - | - |
| *B. sp* | *Ploiarium* | O00B465 | - | EF196016.1 | EF200498.1 | EF202310.1 | **MN619231** | **MN618747** | - | - |
| *B. turkii* | *Ploiarium* | FS1595 | - | EF196073.1 | EF200484.1 | EF202294.1 | **MN619239** | **MN618756** | - | - |
| *B. caespitosum* | *Ploiarium* | P00107296 | **MN639604** | - | - | **MN618950** | - | - | **MN618781** | - |
| *B. clavatum* | *Ploiarium* | P02088221 | - | - | - | - | **MN639653** | - | **MN618785** | - |
| *B. lemuraeoides* | *Ploiarium* | P00097290 | - | - | - | **MN618967** | - | - | **MN618816** | - |
| *B. rubrolabium* | *Ploiarium* | P00097318 | **MN639609** | - | - | - | - | - | **MN618859** | - |
| *B. sp. nov2* | *Ploiarium* | FS2547 | - | **MN618611** | - | **MN618992** | **MN639667** | **MN618741** | **MN618869** | - |
| *B. aggregatum* | *Ploiarium* | FS832 | **MN618462** | EF195951.1 | EF200472.1 | EF202282.1 | **MN619131** | **MN618623** | **MN618763** | **MN619006** |
| *B. ankaizinense* | *Ploiarium* | FS739 | **MN618471** | EF195959.1 | EF200495.1 | EF202306.1 | **MN619140** | **MN618633** | **MN618772** | **MN619016** |
| *B. coccinatum* | *Ploiarium* | FS5961 | **MN618482** | MH822376 | **MN618898** | **MN618953** | **MN639654** | **MN618648** | **MN618786** | **MN619026** |
| *B. coriophorum* | *Ploiarium* | FS649 | **MN618486** | EF195970.1 | EF200474.1 | EF202284.1 | **MN619153** | **MN639616** | **MN618790** | **MN619030** |
| *B. cyclanthum* | *Ploiarium* | K16652, FS5786 | **MN618488** | EF196011.1 | EF200475.1 | EF202285.1 | - | **MN618651** | **MN618791** | **MN619032** |
| *B. divaricatum* | *Ploiarium* | FS4775 | **MN618494** | MH822384 | **MN618904** | **MN618959** | **MN619159** | **MN618655** | **MN618797** | **MN619036** |
| *B. graciliscapum* | *Ploiarium* | FS960 | **MN618495** | EF196010.1 | EF200488.1 | EF202299.1 | **MN619163** | **MN618661** | **MN618800** | **MN619040** |
| *B. henrici* | *Ploiarium* | FS833 | **MN618497** | EF196004.1 | EF200490.1 | EF202301.1 | **MN619165** | **MN618663** | **MN618802** | **MN619042** |
| *B. hirsutiusculum* | *Ploiarium* | FS1926, FS5145 | **MN618498** | EF196064.1 | EF200503.1 | EF202315.1 | **MN619167** | **MN618665** | **MN618803** | **MN619044** |
| *B. hovarum* | *Ploiarium* | FS1397 | - | **MN618588** | - | **MN618961** | **MN639657** | **MN618668** | - | **MN619046** |
| *B. insolitum* | *Ploiarium* | FS648 | **MN618501** | EF196019.1 | EF200478.1 | EF202288.1 | **MN619171** | **MN618675** | **MN618806** | **MN619051** |
| *B. labatii* | *Ploiarium* | FS737 | **MN618506** | EF195948.1 | **MN618907** | EF202296.1 | **MN619174** | **MN618678** | **MN618811** | **MN619055** |
| *B. lancisepalum* | *Ploiarium* | FS1341 | **MN618508** | - | **MN618909** | **MN618966** | **MN619176** | **MN618680** | **MN618813** | **MN619056** |
| *B. lucidum* | *Ploiarium* | FS1040 | **MN618515** | EF195988.1 | EF200493.1 | EF202304.1 | **MN619181** | **MN618687** | **MN618821** | **MN619064** |
| *B. moramanganum* | *Ploiarium* | FS656 | **MN618522** | EF195981.1 | EF200491.1 | EF202302.1 | **MN619188** | **MN618697** | **MN618829** | **MN619073** |
| *B. namoronae* | *Ploiarium* | FS1614 | **MN618526** | MH822404 | **MN618915** | **MN618974** | **MN619192** | **MN618702** | **MN618833** | **MN619076** |
| *B. oenanthum** | *Ploiarium* | FS4418 | **MN618534** | **MN618599** | **MN618916** | **MN618975** | **MN619198** | **MN618707** | **MN618839** | **MN619082** |
| *B. ophiuchus* | *Ploiarium* | FS1480 | **MN618536** | EF195990.1 | EF200500.1 | EF202312.1 | **MN619199** | **MN618708** | **MN618841** | **MN619083** |
| *B. oreodorum* | *Ploiarium* | FS2011 | **MN618537** | EF196043.1 | EF200481.1 | EF202291.1 | **MN619200** | **MN618709** | **MN639674** | **MN619084** |
| *B. ormerodianum* | *Ploiarium* | FS1050 | **MN618538** | MH822407 | **MN618918** | **MN618977** | **MN619201** | **MN639627** | **MN618842** | **MN619085** |
| *B. peyrotii* | *Ploiarium* | FS742 | **MN618544** | EF196051.1 | EF200482.1 | EF202292.1 | **MN619207** | **MN618717** | **MN618848** | **MN619092** |
| *B. platypodum* | *Ploiarium* | K16930 | **MN618546** | EF196052.1 | EF200483.1 | EF202293.1 | - | **MN618718** | **MN618850** | **MN619093** |
| *B. pleiopterum* | *Ploiarium* | S6496 | **MN618547** | **MN618602** | **MN618920** | **MN618980** | **MN619208** | **MN618719** | **MN618851** | **MN619094** |
| *B. protectum* | *Ploiarium* | FS601 | **MN618550** | MH822412 | **MN618923** | **MN618983** | **MN619211** | **MN618721** | **MN618854** | **MN619095** |
| *B. quadrialatum* | *Ploiarium* | FS1080 | **MN639607** | MH822415 | - | **MN618984** | **MN619212** | - | **MN618855** | **MN619097** |
| *B. rubiginosum* | *Ploiarium* | FS1035 | **MN618552** | EF196056.1 | EF200492.1 | EF202303.1 | **MN619213** | **MN618723** | **MN618858** | **MN619099** |
| *B. sp1* | *Ploiarium* | FS4225 | **MN618556** | **MN618605** | - | **MN618987** | **MN619217** | **MN618731** | - | **MN619105** |
| *B. sp11* | *Ploiarium* | FS624 | **MN618558** | EF195995.1 | **MN618924** | **MN618989** | **MN619220** | **MN618733** | **MN618864** | **MN619106** |
| *B. sp. nov9* | *Ploiarium* | FS5489 | - | **MN618613** | - | **MN648668** | **MN619228** | **MN639631** | **MN618872** | **MN619115** |
| *B. sp. nov* | *Ploiarium* | FS4420 | **MN618567** | **MN618615** | **MN618929** | **MN618995** | **MN619229** | **MN618745** | **MN618873** | **MN619116** |
| *B. sp* | *Ploiarium* | FS958 | **MN618564** | EF196009.1 | EF200489.1 | EF202300.1 | **MN619223** | **MN639632** | **MN618874** | **MN619117** |
| *B. subclavatum* | *Ploiarium* | FS907 | **MN618568** | EF196006.1 | EF200476.1 | EF202286.1 | **MN619233** | **MN618749** | **MN618876** | **MN619119** |
| *B. subcrenulatum* | *Ploiarium* | FS5881 | **MN618569** | - | **MN618931** | **MN618998** | **MN619234** | **MN618750** | **MN618877** | **MN619120** |
| *B. therezienii* | *Ploiarium* | FS4426 | **MN618571** | **MN618616** | **MN618932** | **MN618999** | **MN619236** | **MN618752** | **MN639675** | **MN619122** |
| *B. toilliezae* | *Ploiarium* | 792A1398 | **MN618572** | **MN618617** | **MN618933** | **MN619000** | **MN619237** | **MN618753** | **MN618879** | **MN619123** |
| *B. ankaratranum* | *Ploiarium* | MO2599826 | **MN618472** | **MN618580** | **MN618891** | **MN618943** | - | **MN618634** | **MN618773** | **MN639633** |
| *B. callosum* | *Ploiarium* | FS625 | **MN618479** | MH822372 | **MN618897** | **MN618951** | **MN619148** | **MN618643** | **MN618782** | **MN639635** |
| *B. comorianum* | *Ploiarium* | ORCH121223 | **MN618484** | - | **MN618900** | **MN618955** | **MN619152** | **MN618650** | **MN618788** | **MN639637** |
| *B. jackyi* | *Ploiarium* | FS868 | **MN618503** | EF196020.1 | EF200485.1 | EF202295.1 | **MN619172** | **MN639619** | **MN618808** | **MN639639** |
| *B. masoalanum* | *Ploiarium* | FS1082 | **MN618518** | EF195982.1 | EF200480.1 | EF202290.1 | **MN619185** | **MN618693** | **MN618824** | **MN639640** |
| *B. nitens* | *Ploiarium* | FS870 | **MN618530** | EF196034.1 | EF200487.1 | EF202298.1 | **MN619195** | **MN618705** | **MN618837** | **MN639642** |
| *B. prismaticum* | *Ploiarium* | REU9 | **MN618549** | MH822411 | **MN618922** | **MN618982** | **MN619210** | **MN618720** | **MN618853** | **MN639644** |
| *B. sarcorhachis* | *Ploiarium* | FS943 | **MN618554** | EF196008.1 | EF200486.1 | EF202297.1 | **MN619215** | **MN618727** | **MN618861** | **MN639645** |
| *B. petrae* | *Polyradices* | FS2287 | **MN618543** | EF196050.1 | EF200403.1 | - | **MN619206** | **MN618716** | **MN618847** | **MN619091** |
| *B. herbula* | *Trichopus* | Fournel86 | - | **MN618587** | - | - | **MN619166** | **MN618664** | - | - |
| *B. sp. nov* | *Trichopus* | FS4073 | - | **MN618608** | - | **MN618996** | **MN619230** | **MN618746** | - | - |
| *B. boiteaui* | *Trichopus* | S6601 | **MN618477** | **MN618583** | **MN618895** | **MN618947** | **MN639652** | **MN618640** | **MN618778** | - |
| *B. muscicola* | *Trichopus* | FS790 | **MN618525** | EF196033.1 | EF200506.1 | EF202318.1 | **MN619191** | **MN618701** | **MN618832** | - |
| *B. nigriflorum* | *Trichopus* | FS1622 | **MN618529** | EF195993.1 | EF200509.1 | EF202321.1 | **MN619194** | **MN618704** | **MN618836** | - |
| *B. onivense* | *Trichopus* | O00B63 | **MN618535** | **MN618600** | **MN618917** | **MN618976** | - | **MN639626** | **MN618840** | - |
| *B. pantoblepharon* | *Trichopus* | FS957 | **MN618542** | EF196048.1 | EF200507.1 | EF202319.1 | **MN619205** | **MN618713** | **MN618846** | - |
| *B. pleurothallopsis* | *Trichopus* | FS1001 | **MN618548** | EF196053.1 | **MN618921** | **MN618981** | **MN619209** | **MN639628** | **MN618852** | - |
| *B. ciliatilabrum* | *Trichopus* | FS604 | **MN618481** | EF195968.1 | EF200505.1 | EF202317.1 | **MN619150** | **MN618646** | **MN618784** | **MN619024** |
| *B. sp2* | *Trichopus* | FS2659 | **MN618557** | **MN618607** | - | **MN618988** | **MN619218** | **MN639630** | **MN618863** | **MN639647** |
| *B. ambrosia* | *Leopardinae* | - | - | KC568306 | - | - | KF361703 | KF361657 | - | - |
| *B. epiphytum* | *Micranthae* | OR987/09 | - | GQ339693 | GQ339623 | - | - | **MN618656** | - | - |
| *B. affine* | *Sestochilus* | OR209/99 | **MN618460** | EF195916 | **MN618886** | - | KF361704 | KF361658 | **MN618762** | **MN619005** |
| *B. sandersonii* | *Megaclinium* | AFRICA2 | MH822418 | - | **MN618890** | **MN618942** | - | - | **MN618770** | **MN619014** |
| *B. cumingii* | *Cirrhopetalum* | OR181/98 | **MN618487** | EF195923 | EF200404 | EF202322 | - | - | **MN639670** | **MN619031** |
| *B. intertextum* | *Ptiloglossum* | ORCH070350 | **MN618502** | **MN618590** | **MN618905** | **MN618962** | **MN639658** | - | **MN618807** | **MN619052** |
| *B. mayombeense* | *Ptiloglossum* | HBV400/96 | **MN618519** | EF195934 | **MN618912** | **MN618971** | **MN639663** | **MN618694** | **MN618825** | **MN619070** |
| *B. napellii* | *Napellii* | OR76/01 | **MN618527** | GQ339711 | GQ339637 | - | - | **MN618703** | **MN618834** | **MN619077** |
| *B. picturatum* | *Cirrhopetalum* | OR204/98 | **MN618545** | EF195939 | EF200407 | EF202325 | - | - | **MN618849** | **MN639643** |

**Table S2. Features of the four DNA partitions used for phylogenetic inference of Malagasy *Bulbophyllum*.**

|  | **nrITS** | ***Xdh*** | ***PI*** | **5 plastid markers combined** |
| --- | --- | --- | --- | --- |
| Number of taxa | 174 | 143 | 165 | 188 |
| Aligned length (bp) | 743 | 1022 | 458 | 5489 |
| % variable sites | 46.97 | 28.67 | 35.37 | 26.07 |
| Number of constant sites | 394 | 729 | 296 | 4058 |
| Number of potentially parsimony-informative sites | 223 | 120 | 100 | 748 |
| Best fit model (based on AIC) | GTR+F+I+G4 | TPM2u+F+I+G4 | TVM+F+I+G4 | GTR+F+I+G4 |

**Table S3. Testing alternative models of niche transition in Malagasy *Bulbophyllum*.** Results were obtained using bayestraits and logarithmic Bayes Factor (BF) comparisons. There is strong (BF = >6) to very strong (BF = >10) evidence against models that oppose transitions (*q*) from *C* to *B* and from *B* to *A*, respectively (marked in bold).

| **Model** | **Marginal likelihood** | **S.E.** | **log BF** |
| --- | --- | --- | --- |
| Full | -108.342 | +/- 0.009 | 0 |
| *q*_BC_ = 0 | -107.424 | +/- 0.021 | 0.918 |
| *q*_CA_ = 0 | -108.350 | +/- 0.005 | -0.008 |
| *q*_AB_ = 0 | -108.785 | +/- 0.013 | -0.443 |
| *q*_AC_ = 0 | -109.354 | +/- 0.046 | -1.012 |
| ***q*_CB_ = 0** | **-115.238** | **+/- 0.056** | **-6.896** |
| ***q*_BA_ = 0** | **-118.755** | **+/- 0.026** | **-10.413** |

Niche states: *A*, Northwest ‘Sambirano’; *B*, Eastern Lowlands; *C*, Central Highlands.

Abbreviations: S.E., standard error.

**Table S4**. List of 201 accessions representing 192 different *Bulbophyllum* species analysed for photosynthetic pathway (CAM vs. C_3_) via isotope analyses. Of those, 171 species are included in the present phylogeny. Species with δ^13^C values less negative than -20 ‰ are indicated in bold (= CAM species). δ^15^N values are also reported.

| **Species** | **Section** | **Collection** | **in phylogeny** | **δ^13^C (**‰) | **δ^15^N (**‰) |
| --- | --- | --- | --- | --- | --- |
| *B. occlusum* | *Alcistachys* | FS5080 | Yes | -36.99 | -2.74 |
| *B. sp. nov. 1* | *Alcistachys* | FS4391 | Yes | -34.21 | -1.05 |
| *B. variegatum* | *Alcistachys* | FS5975 | Yes | -31.15 | -0.87 |
| *B. sulfureum* | *Alcistachys* | FS1585 | Yes | -29.92 | -0.88 |
| *B. hamelinii* | *Alcistachys* | FS2660 | Yes | -28.53 | 0.50 |
| *B. brevipetalum* | *Alcistachys* | FS2048 | Yes | -28.28 | -2.15 |
| ***B. complanatum*** | ***Bifalcula*** | **FS6986** | **Yes** | **-18.99** | **-2.16** |
| ***B. minutum*** | ***Bifalcula*** | **FS5916** | **Yes** | **-16.57** | **-6.46** |
| ***B. sp. nov. 1*** | ***Bifalcula*** | **FS2707** | **Yes** | **-15.96** | **3.68** |
| ***B. capuroni*** | ***Bifalcula*** | **FS1010** | **Yes** | **-15.61** | **-3.85** |
| ***B. complanatum*** | ***Bifalcula*** | **FS6986** | **Yes** | **-15.35** | **-2.80** |
| ***B. implexum*** | ***Bifalcula*** | **FS6985** | **Yes** | **-15.16** | **-3.09** |
| ***B. complanatum*** | ***Bifalcula*** | **FS5756** | **Yes** | **-13.95** | **0.55** |
| ***B. implexum*** | ***Bifalcula*** | **FS1207** | **Yes** | **-13.71** | **-4.29** |
| ***B. capuroni*** | ***Bifalcula*** | **FS5484** | **Yes** | **-12.15** | **-3.82** |
| ***B. obtusatum*** | ***Calamaria*** | **FS7137** | **Yes** | **-20.20** | **-4.56** |
| ***B. elliotii*** | ***Calamaria*** | **FS5240** | **Yes** | **-18.21** | **-1.57** |
| ***B. incurvum*** | ***Calamaria*** | **FS1081** | **Yes** | **-17.14** | **-2.84** |
| ***B. sp. nov. 1*** | ***Calamaria*** | **FS5246** | **Yes** | **-16.81** | **-3.70** |
| ***B. humblotii*** | ***Calamaria*** | **FS5243** | **Yes** | **-16.80** | **-1.00** |
| ***B. luteobracteatum*** | ***Calamaria*** | **FS5512** | **Yes** | **-16.65** | **-2.81** |
| ***B. erectum*** | ***Calamaria*** | **FS2706** | **Yes** | **-16.58** | **-3.59** |
| ***B. cirrhoglossum*** | ***Calamaria*** | **FS4125** | **Yes** | **-15.62** | **-3.35** |
| ***B. pentastachium*** | ***Calamaria*** | **FS5382** | **Yes** | **-15.58** | **1.16** |
| ***B. malawiense*** | ***Calamaria*** | **OR146_05** | **Yes** | **-14.46** | **-5.25** |
| ***B. pervillei*** | ***Calamaria*** | **FS5444** | **Yes** | **-14.44** | **-4.63** |
| ***B. occultum*** | ***Calamaria*** | **FS5151** | **Yes** | **-14.42** | **-1.96** |
| ***B. histrionicum*** | ***Calamaria*** | **FS6018** | **Yes** | **-14.28** | **-0.01** |
| ***B. maculatum*** | ***Calamaria*** | **FS1133** | **Yes** | **-14.19** | **-1.67** |
| ***B. trifarium*** | ***Calamaria*** | **FS1224** | **Yes** | **-14.10** | **-1.78** |
| ***B. rubrum*** | ***Calamaria*** | **FS5603** | **Yes** | **-14.07** | **7.89** |
| ***B. sambiranense* var. *latibracteatum*** | ***Calamaria*** | **FS5719** | **Yes** | **-13.74** | **-2.04** |
| ***B. pusillum*** | ***Calamaria*** | **FS2281** | **Yes** | **-13.64** | **-4.51** |
| ***B. hildebrandtii*** | ***Calamaria*** | **FS1169** | **Yes** | **-13.49** | **2.09** |
| ***B. bicoloratum*** | ***Calamaria*** | **FS5499** | **Yes** | **-12.98** | **-0.55** |
| ***B. ambongense*** | ***Calamaria*** | **FS6023** | **Yes** | **-12.73** | **0.98** |
| ***B. senghasii*** | ***Calamaria*** | **FS7210** | **Yes** | **-12.62** | **-2.67** |
| ***B. ruginosum*** | ***Calamaria*** | **FS5885** | **Yes** | **-12.29** | **-2.43** |
| ***B. lecouflei*** | ***Calamaria*** | **FS2715** | **Yes** | **-12.09** | **-0.93** |
| ***B. longiflorum*** | ***Cirrhopetalum*** | **FS5990** | **Yes** | **-16.09** | **-5.66** |
| *B. francoisii* | *Elasmatopus* | FS691 | Yes | -37.75 | -3.21 |
| *B. multivaginatum* | *Elasmatopus* | 5164806 | No | -33.17 | -2.61 |
| *B. sp.* | *Elasmatopus* | FS2992 | Yes | -32.99 | -3.78 |
| *B. pandurella* | *Elasmatopus* | FS6641 | Yes | -32.39 | -4.02 |
| *B. amphorimorphum* | *Elasmatopus* | FS2797 | Yes | -32.12 | -2.27 |
| *B. kieneri* | *Elasmatopus* | 3765968 | No | -32.11 | -4.63 |
| *B. oxycalyx* | *Elasmatopus* | FS2959 | Yes | -31.35 | -3.19 |
| *B. aubrevillei* | *Elasmatopus* | FS5200 | Yes | -31.25 | 0.72 |
| *B. analamazoatrae* | *Elasmatopus* | FS773 | Yes | -30.39 | 0.05 |
| *B. sp. nov. 9* | *Elasmatopus* | FS5430 | Yes | -28.27 | -2.34 |
| *B. rauhii_*var*_andranobeense* | *Elasmatopus* | FS1326 | Yes | -28.17 | -2.61 |
| *B. latipetalum* | *Elasmatopus* | FS688 | Yes | -27.31 | -1.90 |
| *B. cardiobulbum* | *Inversiflorum* | FS2971 | Yes | -31.09 | -0.83 |
| *B. uroplatoides* | *Inversiflorum* | FS5055 | Yes | -29.90 | -3.44 |
| ***B. cochinealloides*** | ***Inversiflorum*** | **JH6718** | **Yes** | **-18.58** | **-3.12** |
| *B. alexandrae* | *Kainochilus* | FS5019 | Yes | -33.49 | -1.09 |
| *B. viguieri* | *Kainochilus* | FS2580 | Yes | -30.04 | -6.07 |
| *B. jeanbosseri* | *Kainochilus* | FS7342 | Yes | -29.13 | -2.84 |
| *B. imerinense* | *Kainochilus* | FS2991 | Yes | -29.10 | -2.58 |
| *B. edentatum* | *Kainochilus* | FS6669 | Yes | -29.01 | -0.56 |
| *B. multiligulatum* | *Kainochilus* | FS5612 | Yes | -28.89 | -4.12 |
| *B. cylindrocarpum* | *Kainochilus* | REU7 | Yes | -28.71 | 3.81 |
| *B. reflexiflorum* | *Kainochilus* | 4997682 | No | -28.64 | -4.37 |
| *B. mirificum* | *Kainochilus* | LG4797 | Yes | -27.13 | -4.76 |
| *B. cylindrocarpum* var. *andrngitrense* | *Kainochilus* | FS6857 | Yes | -27.05 | -2.71 |
| *B. sp. nov. 1* | *Kainochilus* | FS5517 | Yes | -25.40 | -3.02 |
| *B. anjozorobeense* | *Kainochilus* | FS1457 | Yes | -23.83 | -4.99 |
| *B. horizontale* | *Kainochilus* | FS7765 | Yes | -21.90 | -2.93 |
| *B. sciaphile* | *Lepiophylax* | FS2785 | Yes | -33.49 | -4.76 |
| *B. jumelleanum* | *Lepiophylax* | FS5791 | Yes | -33.43 | -4.68 |
| *B. calyptropus* | *Lepiophylax* | FS2202 | Yes | -32.95 | -5.86 |
| *B. conchidioides* | *Lepiophylax* | FS1327 | Yes | -29.95 | -2.01 |
| *B. sp_1_Lepiophylax_FS3197* | *Lepiophylax* | FS3197 | Yes | -27.41 | -1.99 |
| *B. sp1* | *Lepiophylax* | FS3197 | Yes | -26.41 | -2.50 |
| *B. bryophytoides* | *Lichenophylax* | FS2595 | Yes | -36.02 | -3.00 |
| *B. debile* | *Lichenophylax* | FS5546 | Yes | -33.94 | -2.98 |
| *B. afzelii* | *Lichenophylax* | FS2526 | Yes | -33.65 | -2.37 |
| *B. mangenotii* | *Lichenophylax* | FS2225 | Yes | -32.54 | 1.77 |
| *B. hapalanthos* | *Lichenophylax* | FS6722 | Yes | -32.22 | 5.00 |
| *B. forsythianum* | *Lichenophylax* | FS1552 | Yes | -31.38 | 0.44 |
| *B. lichenophylax* | *Lichenophylax* | FS5529 | Yes | -31.07 | -0.55 |
| *B. neglectum* | *Lichenophylax* | MO3984 | Yes | -30.33 | -2.50 |
| *B. amparihibense* | *Lichenophylax* | FS4365 | Yes | -30.05 | 1.58 |
| *B. debile* | *Lichenophylax* | FS7661 | Yes | -28.55 | -3.30 |
| *B. sp_nov_FS4002* | *Lichenophylax* | FS4002 | No | -28.39 | -2.94 |
| *B. lakatoense* | *Lichenophylax* | FS3196 | Yes | -27.61 | -1.06 |
| *B. forsythianum* | *Lichenophylax* | FS4740 | Yes | -26.69 | -2.50 |
| *B. melleum* | *Loxosepalum* | MO3759694 | Yes | -35.74 | -1.79 |
| *B. variifolium* | *Loxosepalum* | FS5800 | Yes | -35.48 | -4.42 |
| *B. leandrianum* | *Loxosepalum* | FS3194 | Yes | -35.18 | -5.75 |
| *B. nutans* | *Loxosepalum* | ORCH09049 | Yes | -34.61 | -4.43 |
| *B. minutilabrum* | *Loxosepalum* | 5990279 | No | -33.59 | -4.05 |
| *B. alleizettei* | *Loxosepalum* | FS2335 | Yes | -33.58 | -3.82 |
| *B. leptostachyum* | *Loxosepalum* | 870A981/10 | Yes | -33.06 | -3.67 |
| *B. baronii* | *Loxosepalum* | FS3041 | Yes | -32.45 | -3.19 |
| *B. ambrense* | *Loxosepalum* | FS5886 | Yes | -31.45 | -2.26 |
| *B. subsecundum* | *Loxosepalum* | P00097689 | No | -31.06 | -3.17 |
| *B. tsaratananae* | *Loxosepalum* | 4848627 | No | -30.53 | -3.70 |
| *B. approximatum* | *Loxosepalum* | FS2714 | Yes | -30.46 | -2.02 |
| *B. sp_16* | *Loxosepalum* | FS683 | Yes | -30.36 | -3.85 |
| *B. vakonae* | *Loxosepalum* | FS3141 | Yes | -30.29 | -2.90 |
| *B. florulentum* | *Loxosepalum* | FS1340 | Yes | -30.24 | -1.27 |
| *B. vakonae* | *Loxosepalum* | FS7664 | Yes | -29.70 | -8.07 |
| *B. multiflorum* | *Loxosepalum* | FS7378 | Yes | -29.26 | -7.95 |
| *B. ventriosum* | *Loxosepalum* | P00097607 | No | -28.71 | -3.29 |
| *B. amoenum* | *Loxosepalum* | FS932 | Yes | -28.32 | -3.26 |
| *B. sphaerobulbum* | *Loxosepalum* | 128A479/30 | Yes | -27.87 | -0.08 |
| *B. decaryanum* | *Loxosepalum* | FS949 | Yes | -26.93 | -2.55 |
| *B. lineariligulatum* | *Loxosepalum* | FS1324 | Yes | -26.70 | -1.34 |
| *B. trichochlamys* | *Loxosepalum* | 5990285 | No | -25.98 | -0.90 |
| *B. lyperocephalum* | *Lyperocephalum* | FS6765 | Yes | -32.93 | -6.59 |
| *B. longivaginans* | *Pachychlamys* | FS1402 | Yes | -36.30 | -2.94 |
| *B. vestitum* | *Pachychlamys* | FS5020 | Yes | -34.07 | -1.51 |
| *B. sandrangatense* | *Pachychlamys* | FS5079 | Yes | -33.20 | -2.23 |
| *B. ikongoense* | *Pachychlamys* | FS1514 | Yes | -32.45 | -5.04 |
| *B. sp. nov. 3* | *Pachychlamys* | FS2562 | Yes | -32.08 | -2.92 |
| *B. moratii* | *Pachychlamys* | FS4250 | Yes | -31.75 | -4.03 |
| *B. pachypus* | *Pachychlamys* | FS2918 | Yes | -31.60 | -5.34 |
| *B. macrocarpum* | *Pachychlamys* | P02088165 | Yes | -30.91 | -2.84 |
| *B. molossus* | *Pachychlamys* | FS5789 | Yes | -30.19 | -5.85 |
| *B. liparidioides* | *Pachychlamys* | FS2993 | Yes | -29.65 | -2.22 |
| *B. perrieri* | *Pachychlamys* | P00097716 | No | -29.56 | -3.89 |
| *B. macrocarpum* | *Pachyclamys* | REU5 | Yes | -27.36 | -4.59 |
| *B. discilabium* | *Pantoblepharon* | 279A476/5 | Yes | -31.67 | -2.99 |
| *B. coriophorum* | *Ploiarium* | FS5988 | Yes | -35.52 | -2.30 |
| *B. platypodum* | *Ploiarium* | FS3198 | Yes | -35.01 | -4.71 |
| *B. densum* | *Ploiarium* | P00107318 | Yes | -34.33 | -4.22 |
| *B. sp_nov_2_Ploiarium_FS2547* | *Ploiarium* | FS2547 | Yes | -34.10 | -1.08 |
| *B. rubrigemmum* | *Ploiarium* | FS2941 | No | -34.09 | -1.97 |
| *B. quadrialatum* | *Ploiarium* | ORCH090520 | Yes | -34.06 | -2.15 |
| *B. hovarum* | *Ploiarium* | FS1397 | Yes | -33.83 | -0.99 |
| *B. ormerodianum* | *Ploiarium* | FS1050 | Yes | -33.76 | -2.28 |
| *B. protectum* | *Ploiarium* | FS5212 | Yes | -32.88 | -2.62 |
| *B. andohahelense* | *Ploiarium* | FS2447 | No | -32.83 | -3.95 |
| *B. subcrenulatum* | *Ploiarium* | FS5881 | Yes | -32.79 | -2.01 |
| *B. oenanthum* | *Ploiarium* | FS4418 | Yes | -32.75 | -0.07 |
| *B. insolitum* | *Ploiarium* | FS5875 | Yes | -32.66 | -4.44 |
| *B. subclavatum* | *Ploiarium* | FS2816 | Yes | -32.62 | -8.28 |
| *B. coccinatum* | *Ploiarium* | FS5998 | Yes | -32.55 | -2.52 |
| *B. sp. nov.* | *Ploiarium* | FS4420 | Yes | -32.47 | -2.62 |
| *B. jackyi* | *Ploiarium* | WU0063515 | Yes | -32.39 | -2.89 |
| *B. therezienii* | *Ploiarium* | FS4426 | Yes | -32.30 | -0.31 |
| *B. ankaizinense* | *Ploiarium* | FS1351 | Yes | -32.29 | -3.53 |
| *B. namoronae* | *Ploiarium* | FS6600 | Yes | -32.20 | -0.62 |
| *B. sp. 1* | *Ploiarium* | FS4225 | Yes | -31.88 | -2.06 |
| *B. antongilense* | *Ploiarium* | FS2566 | No | -31.75 | -4.63 |
| *B. divaricatum* | *Ploiarium* | FS1463 | Yes | -31.51 | -2.13 |
| *B. sp. 9* | *Ploiarium* | FS1558 | Yes | -31.45 | 0.19 |
| *B. lucidum* | *Ploiarium* | 910A1731 | Yes | -31.42 | -0.60 |
| *B. ranomafanae* | *Ploiarium* | 4974305 | No | -31.01 | -4.26 |
| *B. hirsutiusculum* | *Ploiarium* | FS1945 | Yes | -30.77 | -0.78 |
| *B. callosum* | *Ploiarium* | FS2332 | Yes | -30.37 | -1.80 |
| *B. nitens* | *Ploiarium* | FS5082 | Yes | -30.37 | -2.93 |
| *B. comorianum* | *Ploiarium* | ORCH121223 | Yes | -30.28 | -4.00 |
| *B. auriflorum* | *Ploiarium* | FS6775 | Yes | -30.25 | -1.14 |
| *B. teretibulbum* | *Ploiarium* | 1881A416 | No | -30.23 | -0.65 |
| *B. lemuraeoides* | *Ploiarium* | P00097290 | Yes | -30.05 | -1.58 |
| *B. masoalanum* | *Ploiarium* | FS5965 | Yes | -30.00 | -0.82 |
| *B. peyrotii* | *Ploiarium* | FS2470 | Yes | -29.54 | -3.35 |
| *B. humbertii* | *Ploiarium* | FS7573 | Yes | -29.50 | -6.17 |
| *B. henricii* | *Ploiarium* | FS5604 | Yes | -29.11 | -2.29 |
| *B. ankaratranum* | *Ploiarium* | 2599826 | Yes | -29.02 | -3.92 |
| *B. lancisepalum* | *Ploiarium* | FS1341 | Yes | -28.91 | -3.77 |
| *B. pleiopterum* | *Ploiarium* | FS6496 | Yes | -28.81 | -4.59 |
| *B. moramanganum* | *Ploiarium* | FS1580 | Yes | -28.76 | -1.81 |
| *B. sp_nov_9_Ploiarium_FS5489* | *Ploiarium* | FS5489 | Yes | -28.73 | -3.09 |
| *B. septatum* | *Ploiarium* | MO5337430 | Yes | -28.69 | -9.93 |
| *B. rutenbergianum* | *Ploiarium* | P00097571 | No | -28.68 | -5.78 |
| *B. oreodorum* | *Ploiarium* | FS2011 | Yes | -28.58 | -6.12 |
| *B. aggregatum* | *Ploiarium* | FS6721 | Yes | -28.57 | 1.02 |
| *B. toilliezae* | *Ploiarium* | 792A1398 | Yes | -28.40 | 0.39 |
| *B. graciliscapum* | *Ploiarium* | FS2146 | Yes | -28.30 | -0.08 |
| *B. sarcorhachis* | *Ploiarium* | FS2882 | Yes | -28.10 | 1.37 |
| *B. paleiferum* | *Ploiarium* | FS5063 | Yes | -27.93 | 0.93 |
| *B. labatii* | *Ploiarium* | FS4257 | Yes | -27.63 | -3.41 |
| *B. duplicatum* | *Ploiarium* | P00097314 | Yes | -27.60 | -5.53 |
| *B. rubrolabium* | *Ploiarium* | P00097318 | Yes | -27.53 | -4.17 |
| *B. abbreviatum* | *Ploiarium* | 4836204 | Yes | -27.42 | -4.94 |
| *B. caespitosum* | *Ploiarium* | P00107296 | Yes | -27.24 | -4.05 |
| *B. prismaticum* | *Ploiarium* | P02088182 | Yes | -26.84 | -5.03 |
| *B. myrmecochilum* | *Ploiarium* | P00214520 | No | -26.19 | 2.37 |
| *B. sp_15* | *Ploiarium* | FS602 | Yes | -26.04 | -1.09 |
| *B. humbertii* | *Ploiarium* | FS794 | Yes | -25.88 | 0.20 |
| *B. Ophiuchus* | *Ploiarium* | FS1480 | Yes | -25.83 | -2.36 |
| *B. turkii* | *Ploiarium* | MO04974304 | Yes | -25.64 | -7.04 |
| *B. crassipetalum* | *Ploiarium* | P00097428 | No | -25.12 | 2.49 |
| *B. clavatum* | *Ploiarium* | P02088221 | Yes | -24.22 | -3.52 |
| ***B. petrae*** | ***Polyradices*** | **FS1331** | **Yes** | **-18.34** | **-3.41** |
| *B. intertextum* | *Ptiloglossum* | ORCH070350 | Yes | -28.85 | 1.50 |
| *B. bryophilum* | *Trichopus* | FS1409 | Yes | -34.28 | -5.15 |
| *B. onivense* | *Trichopus* | FS1964 | Yes | -34.06 | -3.55 |
| *B. boiteaui* | *Trichopus* | FS6601 | Yes | -33.80 | 2.87 |
| *B. muscicola* | *Trichopus* | MO_Razafindasy473 | Yes | -33.64 | -2.50 |
| *B. pantoblepharon* | *Trichopus* | ORCH130064 | Yes | -33.58 | 2.41 |
| *B. sp. 2* | *Trichopus* | FS2659 | Yes | -33.54 | -1.85 |
| *B. pleurothallopsis* | *Trichopus* | FS7371 | Yes | -32.99 | 0.86 |
| *B. bryophilum* | *Trichopus* | FS7049 | Yes | -32.89 | -1.55 |
| *B. nigriflorum* | *Trichopus* | FS1622 | Yes | -31.69 | -4.01 |
| *B. sp_nov* | *Trichopus* | FS4073 | Yes | -27.12 | -6.66 |
| *B. brachystachyum* | # | 5587783 | No | -30.90 | -4.23 |
| *B. leoni* | *#* | P00527405 | No | -30.49 | -4.46 |

**Table S5. Time, palaeo-environmental, and diversity dependent diversification models fitted to the phylogeny of Malagasy *Bulbophyllum*.** Models were fitted to the full tree without outgroup (see **Fig. 1**), its subclade C as well as the tree with subclade C pruned, using rpanda. Number of sampled/total species per (sub)clade are shown in parentheses.

| **Full tree (179/210)** | **LH** | **AICc** | **ΔAICc** | ***λ0*** | ***α*** | ***µ*0** | ***β*** | ***K*** |
| --- | --- | --- | --- | --- | --- | --- | --- | --- |
| **No extinction and constant speciation rate** | **-376.417** | **754.857** | **0** | **0.351** | **–** | **–** | **–** | **–** |
| No extinction and speciation as an exponential function of temperature | -375.856 | 755.780 | 0.923 | 0.206 | 0.038 | – | – | – |
| Linear DD in speciation rate with *K* (= diversity *λ* ~ *µ*) | -374.882 | 755.902 | 1.045 | 0.437 | – | 3.100E-05 | – | 555.615 |
| No extinction and exponential variation in speciation rate | -376.145 | 756.359 | 1.502 | 0.331 | 0.021 | – | – | – |
| Constant speciation and extinction rates | -376.417 | 756.902 | 2.046 | 0.351 | – | 2.481E-08 | – | – |
| Exponential variation in speciation and constant extinction rates | -376.145 | 758.428 | 3.571 | 0.331 | 0.021 | 1.001E-06 | – | – |
| Exponential DD in speciation rate with *K* (= diversity *λ* ~ *µ*) | -376.304 | 758.745 | 3.888 | 0.595 | – | 0.009 | – | Inf |
| Constant speciation rate and exponential variation in extinction rate | -376.417 | 758.971 | 4.115 | 0.351 | – | 5.286E-07 | -0.241 | – |
| Speciation and extinction rates as exponential function of temperature | -375.929 | 760.088 | 5.231 | 0.226 | 0.031 | 4.094E-05 | -0.413 | – |
| Exponential variation in speciation and extinction rates | -376.145 | 760.520 | 5.664 | 0.331 | 0.021 | 5.562E-07 | -0.220 | – |
| **Tree without subclade C (149/171)** | **LH** | **AICc** | **ΔAICc** | ***λ*0** | ***α*** | ***µ*0** | ***β*** | ***K*** |
| **No extinction and constant speciation rate** | **-318.422** | **638.872** | **0** | **0.334** | **–** | **–** | **–** | **–** |
| No extinction and speciation as an exponential function of temperature | -317.686 | 639.453 | 0.581 | 0.173 | 0.047 | – | – | – |
| Linear DD in speciation rate with *K* (= diversity *λ* ~ *µ*) | 316.645 | 639.456 | 0.584 | 0.431 |  | 1.350E-04 | – | 399.868 |
| No extinction and exponential variation in speciation rate | -318.097 | 640.277 | 1.405 | 0.311 | 0.024 | – | – | – |
| Constant speciation and extinction rates | -318.422 | 640.927 | 2.055 | 0.334 | – | 1.688E-07 | – | – |
| Exponential DD in speciation rate with *K* (= diversity λ ~ µ) | -317.684 | 641.533 | 2.661 | 0.713 | – | 0.066 | – | Inf |
| Exponential variation in speciation and constant extinction rates | -318.097 | 642.360 | 3.488 | 0.311 | 0.024 | 4.102E-07 | – | – |
| Constant speciation rate and exponential variation in extinction rate | -318.422 | 643.010 | 4.138 | 0.334 | – | 7.861E-07 | -0.192 | – |
| Speciation and extinction rates as exponential function of temperature | -317.809 | 643.895 | 5.023 | 0.196 | 0.037 | 8.010E-07 | -0.170 | – |
| Exponential variation in speciation and extinction rates | -318.098 | 644.473 | 5.601 | 0.311 | 0.024 | 0.000 | -0.031 | – |
| **Subclade C (30/35)** | **LH** | **AICc** | **ΔAICc** | ***λ*0** | ***α*** | ***µ*0** | ***β*** | ***K*** |
| **No extinction and constant speciation rate** | **-54.695** | **111.533** | **0** | **0.418** | **–** | **–** | **–** | **–** |
| No extinction and exponential variation in speciation rate | -54.682 | 113.809 | 2.276 | 0.433 | -0.017 | – | – | – |
| No extinction and speciation as an exponential function of temperature | -54.684 | 113.813 | 2.281 | 0.529 | -0.018 | – | – | – |
| Constant speciation and extinction rates | -54.695 | 113.834 | 2.302 | 0.418 | – | 2.695E-06 | – | – |
| Linear DD in speciation rate with *K* (= diversity λ ~ µ) | -54.444 | 115.812 | 4.279 | 0.940 |  | 0.320 | – | 41.566 |
| Exponential variation in speciation and constant extinction rates | -54.682 | 116.287 | 4.755 | 0.433 | -0.017 | 1.504E-06 | – | – |
| Constant speciation rate and exponential variation in extinction rate | -54.695 | 116.313 | 4.780 | 0.418 | – | 1.388E-06 | -0.280 | – |
| Exponential DD in speciation rate with *K* (= diversity λ ~ µ) | -54.800 | 116.523 | 4.990 | 1.019 | – | 0.189 | – | 1323.928 |
| Speciation and extinction rates as exponential function of temperature | -54.681 | 118.961 | 7.428 | 0.532 | -0.018 | 0.199 | -1.015 | – |
| Exponential variation in speciation and extinction rates | -54.682 | 118.964 | 7.432 | 0.434 | -0.017 | 9.781E-07 | -0.066 | – |

Abbreviations: AICc, Akaike’s Information Criterion corrected for finite sample size; deltaAICc, difference in the AICc scores between the alternative model and the model with the lowest AICc. *λ*, speciation rate; *µ*, extinction rate; *α*, rate of variation for the exponential speciation model; *β*, rate of variation for the exponential extinction model; LH, log-likelihood; *K*, carrying capacity.

**Table S6. Loadings and *R*^2^ values of bioclim variables (bio1–19) on principal components PC1 and PC2 for locality-climate *datasets 1* and *2* of Malagasy *Bulbophyllum*** (see Methods for details on datasets).

| **Variable** |  | **Dataset 1** | | | |  | **Dataset 2** | | | |
| --- | --- | --- | --- | --- | --- | --- | --- | --- | --- | --- |
|  |  | **PC1** | ***R*^2^** | **PC2** | ***R*^2^** |  | **PC1** | ***R*^2^** | **PC2** | ***R*^2^** |
| Bio1 = annual mean temperature |  | 0.263 | 0.736 | -0.230 | 0.221 |  | 0.262 | 0.756 | 0.220 | 0.204 |
| Bio2 = mean diurnal range |  | -0.219 | 0.509 | -0.186 | 0.145 |  | -0.219 | 0.459 | 0.189 | 0.199 |
| Bio3 = isothermality |  | -0.105 | 0.118 | -0.330 | 0.456 |  | -0.092 | 0.052 | 0.365 | 0.598 |
| Bio4 = temp. seasonality |  | -0.093 | 0.092 | 0.335 | 0.468 |  | -0.106 | 0.128 | -0.360 | 0.449 |
| Bio5 = max. temp. of warmest month |  | 0.228 | 0.552 | -0.261 | 0.285 |  | 0.232 | 0.541 | 0.248 | 0.278 |
| Bio6 = min. temp. of coldest month |  | 0.284 | 0.856 | -0.150 | 0.094 |  | 0.279 | 0.883 | 0.148 | 0.073 |
| Bio7 = temp. annual range |  | -0.218 | 0.506 | -0.057 | 0.014 |  | -0.225 | 0.518 | 0.034 | 0.020 |
| Bio8 = mean temp. of wettest quarter |  | 0.271 | 0.780 | -0.165 | 0.114 |  | 0.269 | 0.804 | 0.149 | 0.095 |
| Bio9 = mean temp. of driest quarter |  | 0.277 | 0.815 | -0.190 | 0.151 |  | 0.273 | 0.830 | 0.179 | 0.143 |
| Bio10 = mean temp. warmest quarter |  | 0.272 | 0.783 | -0.168 | 0.118 |  | 0.270 | 0.810 | 0.151 | 0.096 |
| Bio11 = mean temp. coldest quarter |  | 0.258 | 0.707 | -0.256 | 0.273 |  | 0.256 | 0.725 | 0.251 | 0.256 |
| Bio12 = annual precipitation |  | 0.265 | 0.748 | 0.161 | 0.109 |  | 0.265 | 0.815 | -0.152 | 0.083 |
| Bio13 = prec. of wettest month |  | 0.203 | 0.439 | -0.106 | 0.047 |  | 0.204 | 0.459 | 0.110 | 0.118 |
| Bio14 = prec. of driest month |  | 0.228 | 0.550 | 0.309 | 0.399 |  | 0.225 | 0.556 | -0.313 | 0.406 |
| Bio15 = prec. seasonality |  | -0.192 | 0.389 | -0.339 | 0.482 |  | -0.197 | 0.420 | 0.337 | 0.502 |
| Bio16 = prec. of wettest quarter |  | 0.214 | 0.488 | -0.038 | 0.006 |  | 0.216 | 0.540 | 0.042 | 0.039 |
| Bio17 = prec. of driest quarter |  | 0.235 | 0.587 | 0.296 | 0.367 |  | 0.234 | 0.601 | -0.296 | 0.359 |
| Bio18 = prec. of warmest quarter |  | 0.173 | 0.319 | 0.168 | 0.117 |  | 0.180 | 0.384 | -0.137 | 0.045 |
| Bio19 = prec. of coldest quarter |  | 0.246 | 0.640 | 0.275 | 0.315 |  | 0.245 | 0.667 | -0.271 | 0.290 |

Abbreviations: max., maximum; min. minimum; prec., precipitation; temp., temperature.

**Table S7. bamm diversification rate estimates of the full tree, subclade C as well as the tree without that lineage**. *λ*, speciation rate; *μ*, extinction rate; *r*, net diversification rate (*λ*–*μ*); HPD, highest posterior probability

|  | **mean *λ* (95% HPD)** | **mean *µ* (95% HPD)** | ***r* (95% HPD)** |
| --- | --- | --- | --- |
| Full tree | 0.38 (0.32 – 0.47) | 0.06 (3.51E03 – 0.18) | 0.26 (0.29 – 0.32) |
| Subclade C | 0.36 (0.24 – 0.56) | 0.06 (1.49E03 – 0.24) | 0.18 (0.24 – 0.32) |
| Tree without subclade C | 0.39 (0.32 – 0.48) | 0.06 (3.32E03 – 0.18) | 0.26 (0.30 – 0.32) |

**Supplemental Figures**

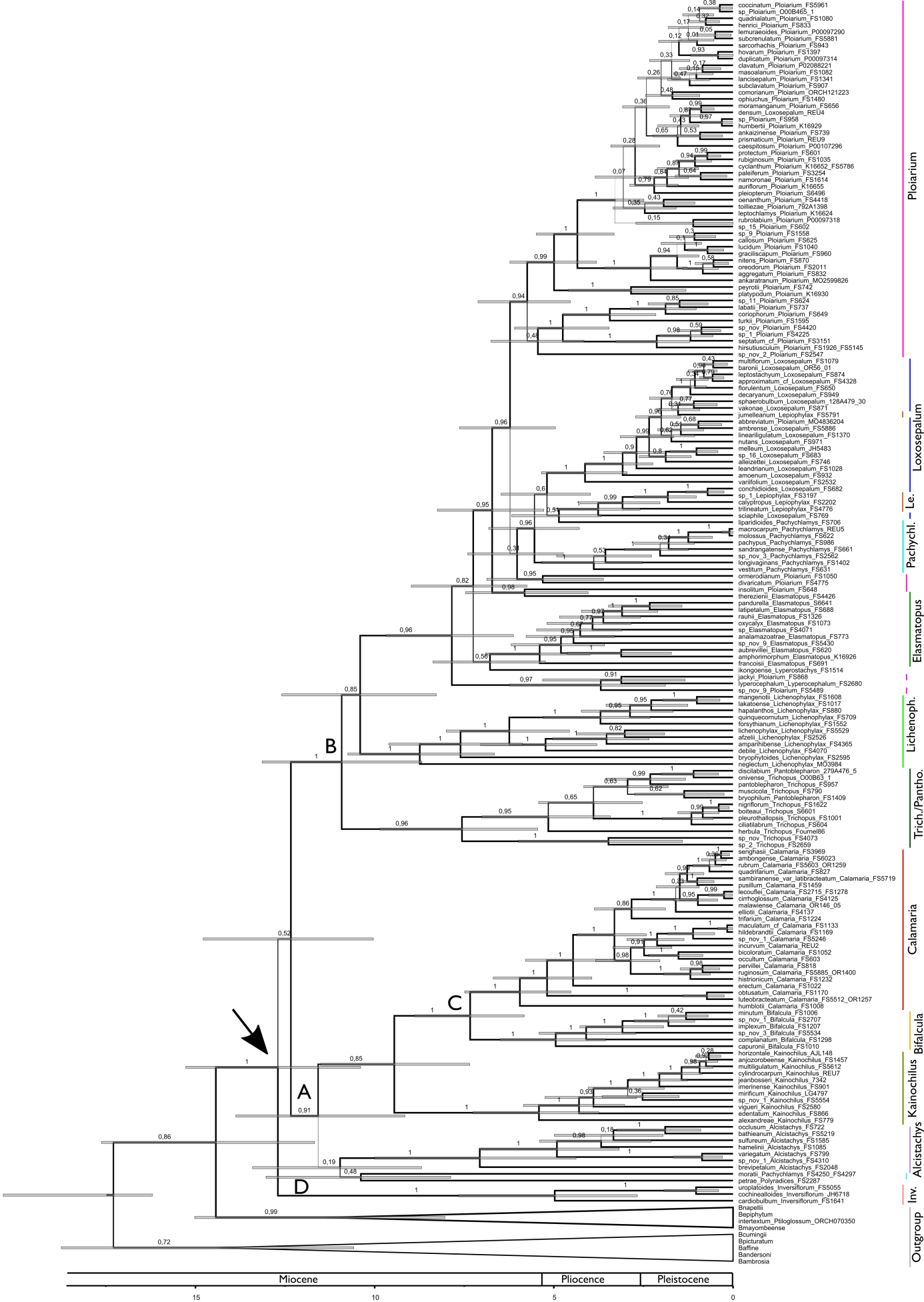

**Fig. S1. beast-derived maximum clade credibility (MCC) chronogram of Malagasy *Bulbophyllum*.** Grey bars indicate 95% highest posterior density (HPD) intervals around mean node ages (in million years ago, MA). Arrow indicates the group’s crown node. Numbers at branches indicate posterior probabilities (PP). Sections are indicated as coloured bars.

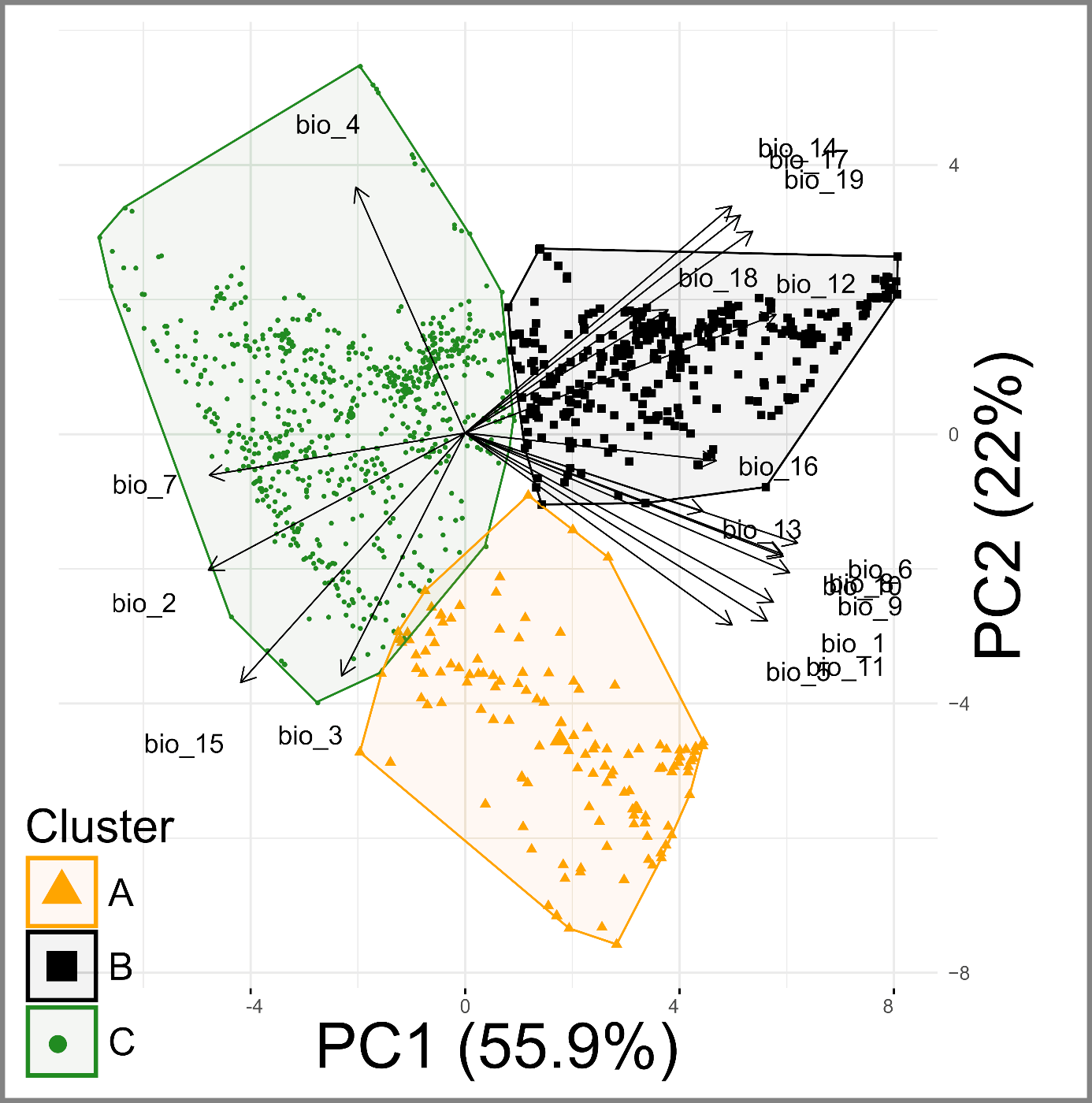

**Fig. S2.** Principal components analysis (PCA) biplot of 1206 occurrence data based on 19 bioclimatic variables (bio1–19) (Note: the factor loadings do not correspond to the x or y axes; see Supplemental Table S6 for component loadings). Each data point is coloured and symbolized according to its cluster membership as defined by *k-means* clustering.
